# The *Arabidopsis* splicing factor PORCUPINE/SmE1 orchestrates temperature-dependent root development via auxin homeostasis maintenance

**DOI:** 10.1101/2024.06.05.597578

**Authors:** Nabila El Arbi, Sarah Muniz Nardeli, Jan Šimura, Karin Ljung, Markus Schmid

**Affiliations:** Umeå Plant Science Centre, Department of Plant Physiology, Umeå University, SE-901 87 Umeå, Sweden; Department of Plant Biology, Linnean Center for Plant Biology, Swedish University of Agricultural Sciences, S-75007 Uppsala, Sweden; Umeå Plant Science Centre, Department of Forest Genetics and Plant Physiology, Swedish University of Agricultural Sciences, SE-901 83 Umeå, Sweden

**Author notes:** **Author responsible for distribution of materials:** Markus Schmid.

**Keywords:** *Arabidopsis thaliana*, alternative RNA splicing, temperature signaling, auxin signaling, root development, SmE, root apical meristem

## Abstract

- Appropriate abiotic stress response is pivotal for plant survival and makes use of multiple signaling molecules and phytohormones to achieve specific and fast molecular adjustments. A multitude of studies has highlighted the role of alternative splicing in response to abiotic stress, including temperature, emphasizing the role of transcriptional regulation for stress response. Here we investigated the role of the core splicing factor *PORCUPINE* (*PCP*) on temperature-dependent root development.
- We used marker lines and transcriptomic analyses to study the expression profiles of meristematic regulators and mitotic markers, and chemical treatments, as well as root hormone profiling to assess the effect of auxin signaling.
- The loss of *PCP* significantly alters RAM architecture in a temperature-dependent manner. Our results indicate that PCP modulates the expression of central meristematic regulators and is required to maintain appropriate levels of auxin in the RAM.
- We conclude that alternative pre-mRNA splicing is sensitive to moderate temperature fluctuations and contributes to root meristem maintenance, possibly through the regulation of phytohormone homeostasis and meristematic activity.

## Introduction

Plants display remarkable phenotypic plasticity, which they achieve by continuously assessing their environment (Guo *et al*., 2018; Lamers *et al*., 2020). Generally, abiotic stresses induce a signaling cascade involving small signaling molecules, such as reactive oxygen species (ROS), and various phytohormones, such as auxin (Lamers *et al*., 2020; Danve *et al*., 2021). It is noteworthy, that while different stresses utilize a common set of signaling components, plants achieve sophisticated and stress-specific transcriptional and physiological responses (Lamers *et al*., 2020). To date, it remains an open question in plant physiology how these stress-specific responses are attained.

Root development and architecture have a strong impact on overall plant physiology, growth rate and stress resistance (Jung & McCouch, 2013; Kuriakose & Silvester, 2016; González-García *et al*., 2023). Yet, many molecular processes governing root growth and patterning, particularly in response to environmental cues, remain unknown (Kuriakose & Silvester, 2016; Motte *et al*., 2019). Briefly, root development and growth are governed by the rate of cell proliferation and elongation (Greb & Lohmann, 2016; Kuriakose & Silvester, 2016; Motte *et al*., 2019). The root apical meristem (RAM) is thus pivotal for root growth, since it harbors the quiescent center (QC), comprising undifferentiated cells, and pluripotent stem cells, which undergo asymmetric cell division to generate daughter cells (Kuriakose & Silvester, 2016). WUSCHEL-LIKE HOMEOBOX5 (WOX5) is an essential transcription factor for the regulation of root development. It is expressed in the QC (Sarkar *et al*., 2007) where it suppresses the expression of *CYCD3;3* and *CYCD1;1*, thus inhibiting cell proliferation (Forzani *et al*., 2014; Motte *et al*., 2019). Root organization can be divided into radial patterning, encompassing vascular tissue, endodermis, cortex and epidermis, and tangential patterning, specifically the differentiation of epidermal cells into trichoblasts (hair-bearing cells) and atrichoblasts (non-hair-bearing cells) (Kuriakose & Silvester, 2016). *Arabidopsis thaliana* (*A. thaliana*) has a type III root hair pattern, where trichoblasts are in contact with two underlying cortical cells, while atrichoblasts are only in contact with one cortical cell, thus creating an organized and repetitive pattern of root hair and non-root hair cells (Salazar-Henao *et al*., 2016). It has been shown that all the above-mentioned processes are governed by auxin and maintaining a stable auxin maximum in the QC and a gradient along the root axis is crucial for the regulation of cell division and expansion (Kuriakose & Silvester, 2016; Zluhan-Martínez *et al*., 2021). Auxin is believed to act upstream of the major regulators of stem cell activity, and auxin concentration can attenuate signaling cascades by modulating gene expression (Kuriakose & Silvester, 2016). Furthermore, research on the role of auxin in root development highlighted that various environmental cues and hormonal signals converge onto auxin signaling (Olatunji *et al*., 2017; Motte *et al*., 2019). It appears that auxin biosynthesis and homeostasis contribute to environmental adaptation. Besides the free, biologically active form of auxin, IAA, it can be found in three different conjugated forms: (a) sugar esters, (b) amide conjugations to amino acids, and (c) amide conjugations to peptides or proteins (Ruiz Rosquete *et al*., 2012; Casanova-Sáez *et al*., 2021). Until recently, it was postulated that among the amino acid conjugations all but two, IAA-Asp and IAA-Glu, were reversible (Ludwig-Müller, 2011; Ruiz Rosquete *et al*., 2012; Korasick *et al*., 2013; Casanova-Sáez *et al*., 2021). However, a new model has been proposed, suggesting that IAA-Asp and IAA-Glu are also reversible conjugations, but can be subject to oxidation, which ultimately leads to auxin inactivation (Hayashi *et al*., 2021; Luo *et al*., 2023). The conjugation of auxin to amino acids depends on the activity of Gretchen Hagen 3 (GH3) family proteins, which also contribute significantly to the regulation of plant stress responses (Ruiz Rosquete *et al*., 2012; Wojtaczka *et al*., 2022; Casanova-Sáez *et al*., 2022; Luo *et al*., 2023).

Low or high temperature are important plant stressors and play a central role in governing developmental processes (Quint *et al*., 2016; Guo *et al*., 2018; Lamers *et al*., 2020; De Smet *et al*., 2021; Penfield *et al*., 2021; Zhu *et al*., 2022). In the case of *A. thaliana*, chilling stress is generally experienced between 0-14°C, wherein exposure to 0-5°C induces cold acclimation, and temperatures <0°C induce freezing stress (Praat *et al*., 2021). However, natural variations between accessions can have a major impact on the temperature sensitivity of *A. thaliana* (Hannah *et al*., 2006; Hernandez *et al*., 2023). Interestingly, it has been reported that at the transcriptome level, temperature response in roots is remarkably different from that in shoots (Bellstaedt *et al*., 2019; Lamers *et al*., 2020). In recent years, several studies have emphasized the role of alternative splicing (AS) in response to temperature (Calixto *et al*., 2018; Neumann *et al*., 2020; Dikaya *et al*., 2021), underlining the necessity of transcriptomic adjustments to temperature cues. Several studies have also highlighted the importance of AS to the cold response in plants (Reddy *et al*., 2013; Staiger & Brown, 2013; Laloum *et al*., 2018; Calixto *et al*., 2018; Capovilla *et al*., 2018). A potential new candidate, which connects plant development, cold temperature response and AS is *PORCUPINE* (*PCP*/*AT2G18740*), which encodes the *A. thaliana* SmE1/SmEb protein (Capovilla *et al*., 2018; Huertas *et al*., 2019; Wang *et al*., 2022), which is an essential part of the spliceosomal SM-ring (Matera & Wang, 2014). *PCP* is induced by cold temperatures and osmotic stress (Cao *et al*., 2011; Capovilla *et al*., 2018; Huertas *et al*., 2019). The *pcp-1* mutant exhibits strong developmental defects when grown at cold temperatures (Capovilla *et al*., 2018; Huertas *et al*., 2019; Wang *et al*., 2022), and a short root phenotype in response to salt stress (Willems *et al*., 2023; Hong *et al*., 2023). Furthermore, *pcp-1* mutants are less sensitive to ROS induced cell damage (Willems *et al*., 2023). However, while these studies provide important insights into the effects of *pcp-1* on transcriptomic changes, or shoot development, the role of PCP in regulating root development has been largely neglected. Here we show that PCP is key to temperature-dependent RAM maintenance by modulating the expression of central meristematic regulators, such as *WOX5*, and by preserving appropriate levels of auxin.

## Materials and Methods

### Plants and growth conditions

A list of all plants and accession numbers can be found in Table S1.

For root growth assays, hormone profiling, and microscopy, seeds were surface sterilized in a solution comprising 70% EtOH, 10% sodium hypochlorite (VWR, 7681-52-9), and 0.01% Triton™ X-100 (Merck, T8787), washed twice with absolute EtOH and dried in a sterile laminar flow cabinet. Sterilized seeds were placed on ½ strength Murashige and Skoog (MS) (Duchefa Biochemie, M0222) growth medium, containing 1.6% plant agar (Duchefa Biochemie, P1001) and 0.5 g/L MES buffer (Duchefa Biochemie, M1503), pH=5.7 using KOH. The seeds were stratified at 4°C in darkness for 48 hours, transferred to Percival growth chambers equipped with full-spectrum white light LEDs, and cultivated at 16°C, 23°C, or 27°C (± 0.5°C), under LD conditions (16h light, 8h dark), with a relative humidity of 65%.

For crossings and seed propagation, seeds were stratified in 0.1% agarose (VWR, 35-1020) solution at 4°C in darkness for 48 hours, then transferred to soil (3:1 soil:vermiculite (Sibelco Europe)). Plants were fertilized once with Rika (SW Horto, Horto Liquid Rika S, 7-5-1) (dilution factor 1:100) and treated with NEMAblom (BioNema AB). Plants were grown in Percival growth chambers equipped with full-spectrum white light LEDs and cultivated at 16°C or 23°C (± 0.5°C), under LD conditions (16h light, 8h dark), with a humidity of 65%.

Hormone treatments with auxinole (MedChemExpress, HY-111444) and IAA (Merck/Sigma-Aldrich, I2886) were carried out on ½MS plates. Each compound was dissolved in DMSO (Merck, D8418) to create stock solutions of 50 mM auxinole and 100 mM IAA. Auxinole was directly added to the growth medium to reach the desired concentrations, while IAA was first further diluted in sterile water (1:1000 ≙ 100 µM) and then added to the growth medium to reach the desired concentrations. Diluted DMSO was used in control plates. IAA-containing plates were wrapped in yellow plastic sheets to prevent light-mediated IAA degradation.

### DNA extraction and plant genotyping

For *pcp-1* genotyping, the Phire Plant Direct PCR Master Mix (Thermo Scientific™, F160L) was used according to manufacturer’s instructions. Briefly, a small pipette tip was used to take a leaf punch, which was then homogenized in 20 µL of dilution buffer. 1 µL of this mixture was used in a PCR mix containing 12.5 µL of Phire Plant Direct PCR Master Mix (2X), 10.5 µL nuclease-free water and 0.5 µL of each oligo. The oligo annealing temperature was 55°C. The oligo sequences were as follows:

PCP-forward (LP): CTCCGATTCACCAGACTTGAG, PCP-reverse (RP): GCCGAAGAGAATGACACAATC, T-DNA (LBb1.3): ATTTTGCCGATTTCGGAAC

### Microscopy

For marker-line, RAM, and stele microscopy we followed the published ClearSee protocol (Kurihara *et al*., 2015), with the following changes: Seedlings were fixed for 1h in 4% PFA + 0.01% Triton™ X-100 and cleared in ClearSee solution overnight. For the PlaCCI -marker line fixation time was shortened to 10 min. After overnight clearing, all seedlings, apart from PlaCCI, were counterstained with SR2200 cell wall staining (Musielak *et al*., 2016) following the protocol established by Tofanelli et al., 2019.

PIN protein immunolocalization was performed using an InsituProVsi (Intavis Bioanalytical Instruments AG) as previously described (Sauer *et al*., 2006; Doyle *et al*., 2015). Seedlings were either grown for 5 days at 23°C (± 0.5°C), or 10 days at 16°C (± 0.5°C) and then sampled. This was made necessary due to sample size restrictions of the InsituProVSI Robot wells. The full protocol for the immunolocalization with the InsituProVSI Robot can be found in Methods S1. The samples were treated with the following antibodies: anti-PIN1 (sheep) (NASC ID: N782246), anti-PIN2 (rabbit) (Abas *et al*., 2006) (from Dr. Siamsa Doyle), anti-PIN7 (rabbit)(Doyle *et al*., 2019) (from Dr. Siamsa Doyle), anti-sheep CY3 (Jackson ImmunoResearch, 713-165-003) and anti-rabbit CY3 (Jackson ImmunoResearch, 111-165-003) (Table S2).

Confocal microscopy was performed using a Zeiss LSM780 CLSM with inverted stand. For RAM phenotyping, marker-line analysis, and immunolocalization, a 40x/1.2 water immersion objective was used. For cell counting of the proximal meristem and PlaCCI visualization, a 25x multi-immersion objective was used. The wavelengths which were used for each marker can be found in Table S3.

For root hair phenotyping, seedlings were imaged at a fluorescence stereomicroscope Leica M205 FA, which was upgraded to the THUNDER Imager Model Organism during this study.

Image analysis was done using FIJI (https://imagej.net/software/fiji/). For better visibility, the contrast of some images has been adjusted. In this case, all images of one panel were adjusted identically.

### Hormone profiling

Root tissue was sampled by cutting the roots below the hypocotyl using a razor blade and immediately snap-frozen. The frozen root tissue was homogenized in liquid nitrogen using mortar and pestle. 5 biological replicates per genotype were sampled.

Samples were extracted, purified, and analyzed according to the method described in Šimura *et al*., 2018 with the inclusion of compounds IAA-Glc and oxIAA-Glc (MRMs, 176,1>130,1 and 192,1>146,1, respectively) and their ^13^C_6_-analogs were used for precise quantification (MRMs, 182,1>130,1 and 198,2>152,1) as described in Pênčík *et al*., 2018. Briefly, approx. 10 mg of frozen material per sample was homogenized and extracted in 1 mL of ice-cold 50% aqueous acetonitrile (v/v) with the mixture of ^13^C- or deuterium-labeled internal standards using a bead mill (27 Hz, 10 min, 4°C; MixerMill, Retsch GmbH). After centrifugation (14 000 RPM, 15 min, 4°C), the supernatant was purified as follows: A solid-phase extraction column Oasis HLB (30 mg 1 mL, Waters Inc.) was conditioned with 1 mL of 100% methanol and 1 mL of deionized water (Milli-Q, Merck Millipore). After the conditioning steps each sample was loaded on SPE column and flow-through fraction was collected with the elution fraction, 1 mL 30% aqueous acetonitrile (v/v). Samples were evaporated to dryness using a speed vac vacuum concentrator (SpeedVac SPD111V, Thermo Scientific). Prior to LC-MS analysis, samples were dissolved in 40 µL of 20% acetonitrile (v/v) and transferred to insert-equipped vials, 20 µL were injected onto the column. Mass spectrometry analysis of targeted compounds was performed by an UHPLC-ESI-MS/MS system comprising of a 1290 Infinity Binary LC System coupled to a 6490 Triple Quad LC/MS System with Jet Stream and Dual Ion Funnel technologies (Agilent Technologies). The quantification was carried out in Agilent MassHunter Workstation Software Quantitative (Agilent Technologies). Primary data and statistical analysis can be found in Tables S4-S5.

### Statistical Analysis

For root length measurement, plates were scanned at an EPSON Expression 12000XL scanner equipped with SilverFast®8 software. Root lengths were measured using the Simple Neurite Tracer found in FIJI. Confocal images were imported into FIJI and cell numbers were counted using the inbuilt cell counter. As an approximation to meristem size, we counted the number of cells in the proximal meristem of the seedlings (number of cortical cells from QC to elongation zone, where cells double in size) (Verbelen *et al*., 2006; Pavelescu *et al*., 2018). To account for different root lengths, stele cells were counted in the differentiated root (two visible protoxylem strands) (Kondo *et al*., 2014). Cell counts, root lengths and hormone quantifications were statistically analyzed using GraphPad Prism 10.2.1. All GraphPad analysis sheets can be found in the supplementary Tables S6-S7.

### RNA extraction, strand-specific RNA sequencing and data analysis

Root tissue was sampled at ZT=6 (Zeitgeber Time, beginning of day is ZT=0) and snap-frozen in liquid nitrogen. The tissue was then homogenized using a bead mill (30 hz, 1 min) and RNA was extracted using TRIzol (Invitrogen™, 15596026) following manufacturer’s instructions. Subsequently, RNA was treated with DNaseI (Invitrogen™, EN0521) following manufacturer’s instructions and sent for strand-specific RNA sequencing at BIOMARKER TECHNOLOGIES (BMK) GmbH, Germany. The raw sequencing data are available at the European Nucleotide Archive (ENA) at EMBL-EBI under the accession number PRJEBxxxx.

Data pre-processing included quality control where the raw sequence data were assessed (FastQC v0.10.1) and ribosomal RNA (SortMeRNA v2.1b) (Kopylova *et al*., 2012) and adaptor sequences (Trimmomatic v0.32) (Bolger *et al*., 2014) were removed, followed by another quality control step to ensure that no technical artifacts were introduced during data pre-processing. mRNA sequences were aligned with Salmon (v0.14.2) (Patro *et al*., 2017) to the *A. thaliana* Reference Transcript Dataset 2 (Zhang *et al*., 2017).

Post processing and analysis of the RNA sequencing data were performed using the 3D RNA sequencing app (Guo *et al*., 2021) (https://3drnaseq.hutton.ac.uk/app_direct/3DRNAseq/). Batch effect removal in the auxinole dataset was performed using the inbuilt batch effect removal (RUVr from the RUVseq package in R). We chose TMM (weighted trimmed mean of M-values) as data normalization method and the output data was filtered for an Absolute log_2_FC of 2 or higher. For further clustering and network inference analysis, the Dashboard for the Inference and Analysis of Networks from Expression data (DIANE) (Cassan *et al*., 2021) was used (https://diane.bpmp.inrae.fr/). Results from the 3DApp and DIANE can be found in Table S8.

## Results

### The loss of *PORCUPINE* causes severe defects of root development

Huertas *et al*., 2019 reported that *pcp-1* seedlings grown at 20°C display shortened roots and increased root hair density. We performed a detailed phenotypic analysis of *pcp-1* root development to gain insight into the molecular mechanisms underlying this phenotype. We observed that *pcp-1* roots were always shorter than those of Col-0, which is in line with the observations from Huertas *et al*., 2019 (Fig. 1**a**-**b**). At the microscopic level the RAM architecture of *pcp-1* was perturbed at 23°C, particularly in the stele (Fig. 1**c**). *pcp-1* seedlings grown at 16°C showed a severe disruption of RAM organization and early differentiation, demonstrated by the early onset of root hairs. Interestingly, 2 days shift to 16°C was sufficient to induce similar phenotypic alterations. The number of cells in the proximal meristem was lower in *pcp-1* at 23°C and was reduced even more at 16°C (Fig. 1**d**). Due to the defects in the stele we wanted to investigate whether stele cell numbers were also affected by the loss of *PCP* and low temperature. While stele cell numbers were significantly reduced in *pcp-1* in comparison to Col-0 (Fig. S1 **a**-**b**), there was no difference due to temperature, suggesting that this may be a genotype-specific phenotype. To test this hypothesis, we grew seedlings at slightly elevated temperatures (27°C). Interestingly, we observed a near full suppression of the *pcp-1* root phenotype at 27°C, with no differences in cell numbers in the proximal meristem or stele compared to Col-0 (Fig. S1).

**Figure 1.**
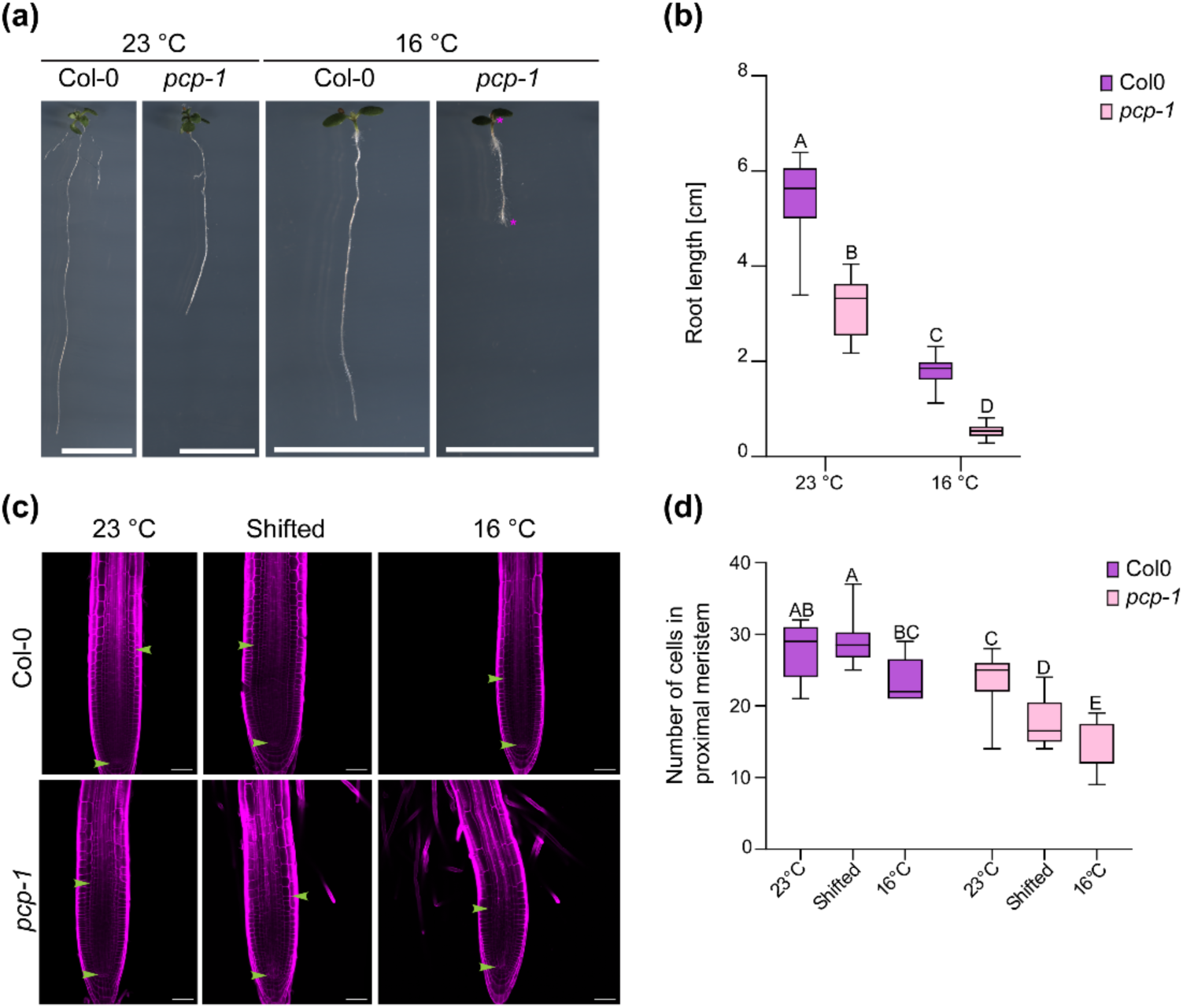
The loss of *pcp-1* causes severe root development defects at 16°C. (a-b) *pcp-1* mutants grown at 16°C exhibit shortened, and hairy roots compared to Col-0. Magenta asterisks indicate the lack of leaf primordia and hairy root tip. Scale bars: 1 cm. (c-d) RAM architecture and the number of cells in the proximal meristem is significantly lower in *pcp-1*. Bottom green arrows: position of QC, top green arrows: beginning of cell elongation. Scale bars: 50 µm. Box plots are min to max. Statistical test: Two-way ANOVA with posthoc Tukey analysis.

Taking a closer look at the RAM organization of *pcp-1* grown at 16°C, we noted that in many seedlings it was challenging to determine the exact position of the QC. Due to the shortened meristem and early differentiation, we thus hypothesized that the meristem could be depleted, or that QC identity could be lost. To investigate this, we crossed *pcp-1* with the established *pWOX5::ER-GFP* marker (Sarkar *et al*., 2007). In the control line, GFP signal was confined to the QC irrespective of growth temperature. Unexpectedly, *pcp-1* mutants carrying the marker and grown at 16°C displayed a widened expression domain, which stretched several cell layers into the stele (Fig. 2**a**). Since WOX5 inhibits cell division in the QC, we were curious to further investigate the number of dividing cells in the root meristem of *pcp-1*. To this end, we employed the established PlaCCI (Plant Cell Cycle Indicator), which combines three fluorescent markers which are expressed during different stages of the cell cycle (Desvoyes *et al*., 2020). We were particularly interested in the *pCycB1;1*::*N-CycB1;1-YFP* expression, since it marks the transition of G2 to M-phase in the cell cycle and thus active cell division. In the control, we observed a significant increase in CycB1;1 positive cells in seedlings grown at 16°C compared to 23°C. In *pcp-1* mutants crossed with the PlaCCI, we observed a slightly reduced number of CycB1;1 expressing cells at 23°C, and a further reduction at 16°C (Fig. 2**b**-**c**). *pCDT1a*::*CDT1a-eCFP* expression is indicative of highly proliferative cells or cells undergoing endocycle (Lasok *et al*., 2023). In the control, we observed CDT1a expression throughout the meristem, which was concentrated around the initials and in the stele. In the mutant, we observed overall weaker, and more diffuse CDT1a expression, and at 16°C, the expression boundary was located closer to the root tip (Fig. 2**b**). Finally, *pHTR13::HTR13-mCherry* expression, and thus histone 3.1 deposition, is associated with proliferation potential (Otero *et al*., 2016). HTR13 expression in *pcp-1* mutants grown at 16°C was markedly reduced in length, displaying a shortened expression domain compared to the control, in which the expression domain extended throughout the observable root (Fig. 2**b**).

**Figure 2.**
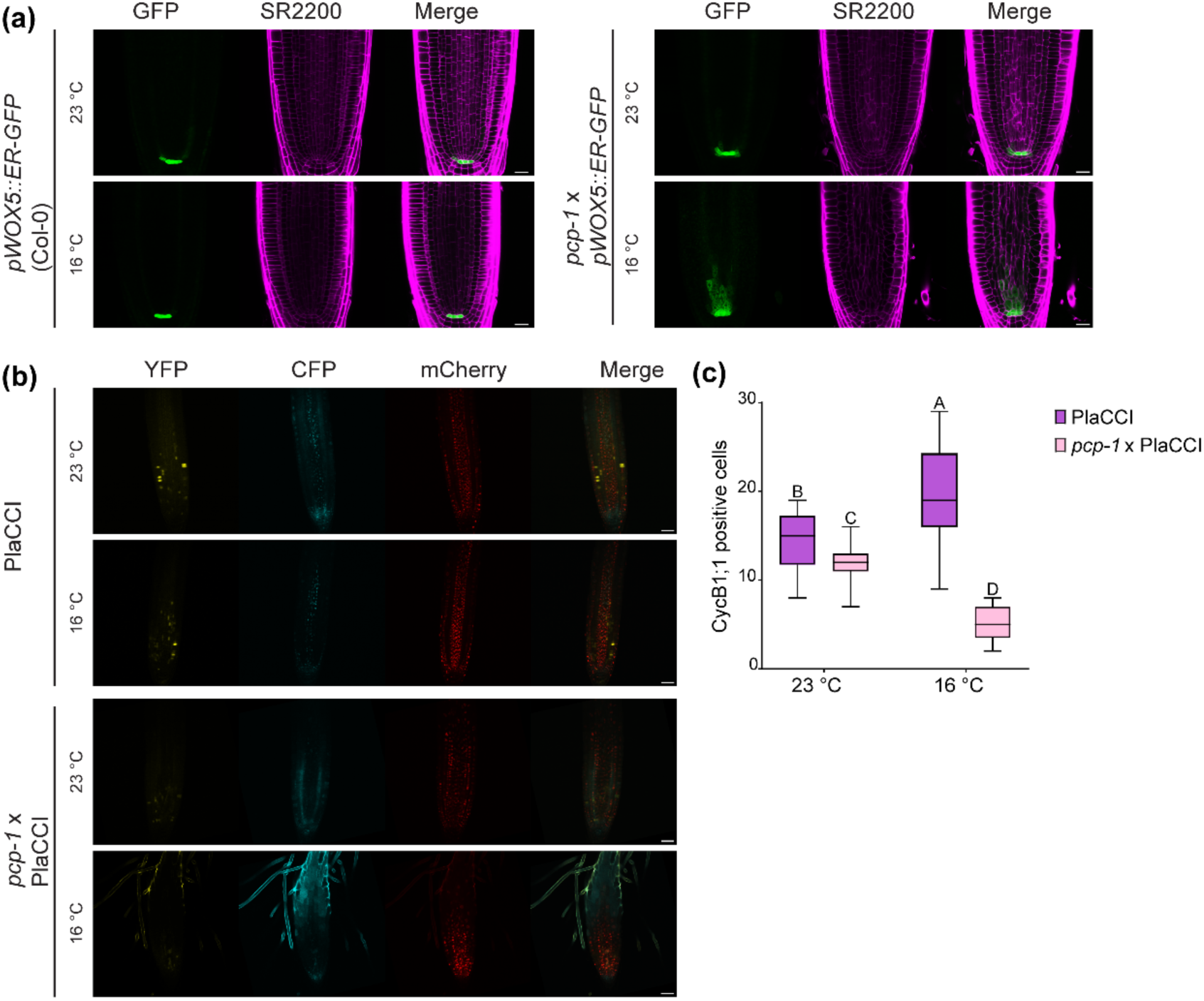
Ectopic *WOX5* expression and reduced number of mitotic cells in *pcp-1* mutants at 16°C. (a) *pWOX5::ER-GFP* expression is restricted to the QC in Col-0 at both 23°C and 16°C. *pcp-1* mutants show ectopic expression of GFP in the stele at 16°C. Scale bars: 20 µm (b) Analysis of the PlaCCI reveals lower expression of CycB1;1 (YFP) and CDT1a (CFP) in *pcp-1* mutants, as well as shortened expression area of HTR13 (mCherry). Scale bars: 50 µm. (c) Quantification of CycB1;1 positive cells. Box plots are min to max. Statistical test: Two-way ANOVA with posthoc Tukey analysis.

Taken together, these results indicate that PCP is essential for RAM maintenance in a temperature-dependent manner. The loss of *PCP* causes an ectopic expression of the QC marker WOX5 at 16°C, correlated with a decrease in cell division, shown by reduced expression of mitotic cell markers, and a shortened meristem.

### *pcp-1* mutants exhibit misspecification of (a)trichoblast cells and elevated endogenous IAA

Our previous results showed that the loss of *PCP* caused pronounced changes in RAM architecture, as well as early onset of root hairs at low temperatures. These observations raised two questions: Firstly, whether root hair cells in *pcp-1* were correctly positioned, and secondly, whether this phenotype was connected to the mis-regulation of endogenous auxin, a known regulator of root hair development (Vissenberg *et al*., 2020). To address these questions, we first crossed *pcp-1* with the *pGL2::SAND* atrichoblast and the *pEXP7::SAND* trichoblast marker lines, derived from the SWELLINE marker set (Marquès-Bueno *et al*., 2016). We observed that the YFP signal in *pGL2::SAND* control lines was restricted to cell files 2-3 cells wide, which were organized in lanes, irrespective of growth temperature. In *pcp-1* crossed with the *pGL2::SAND* marker, we observed that already at 23°C these cell files were not as structured as in the control, displaying emergence or loss of YFP signal within presumed (a)trichoblast lanes, respectively (Fig. 3**a**). Similarly, at 16°C the expression pattern seemed fully disrupted, and atrichoblast lane width appeared to be reduced to 1 cell (Fig. 3**a**). We observed a similar perturbation of the root hair patterning in the *pEXP7::SAND* marker (Fig. 3b). In the control, YFP signal was restricted to epidermal cells touching two underlying cortical cells. In *pcp-1* crossed with *pEXP7::SAND* we observed signal in a few cells only touching one cortical at 23°C, but the correct pattern was mostly preserved. At 16°C, however, the patterning was fully disrupted and YFP signal was found in multiple adjacent epidermal cells, regardless of the number of underlying cortical cells.

**Figure 3.**
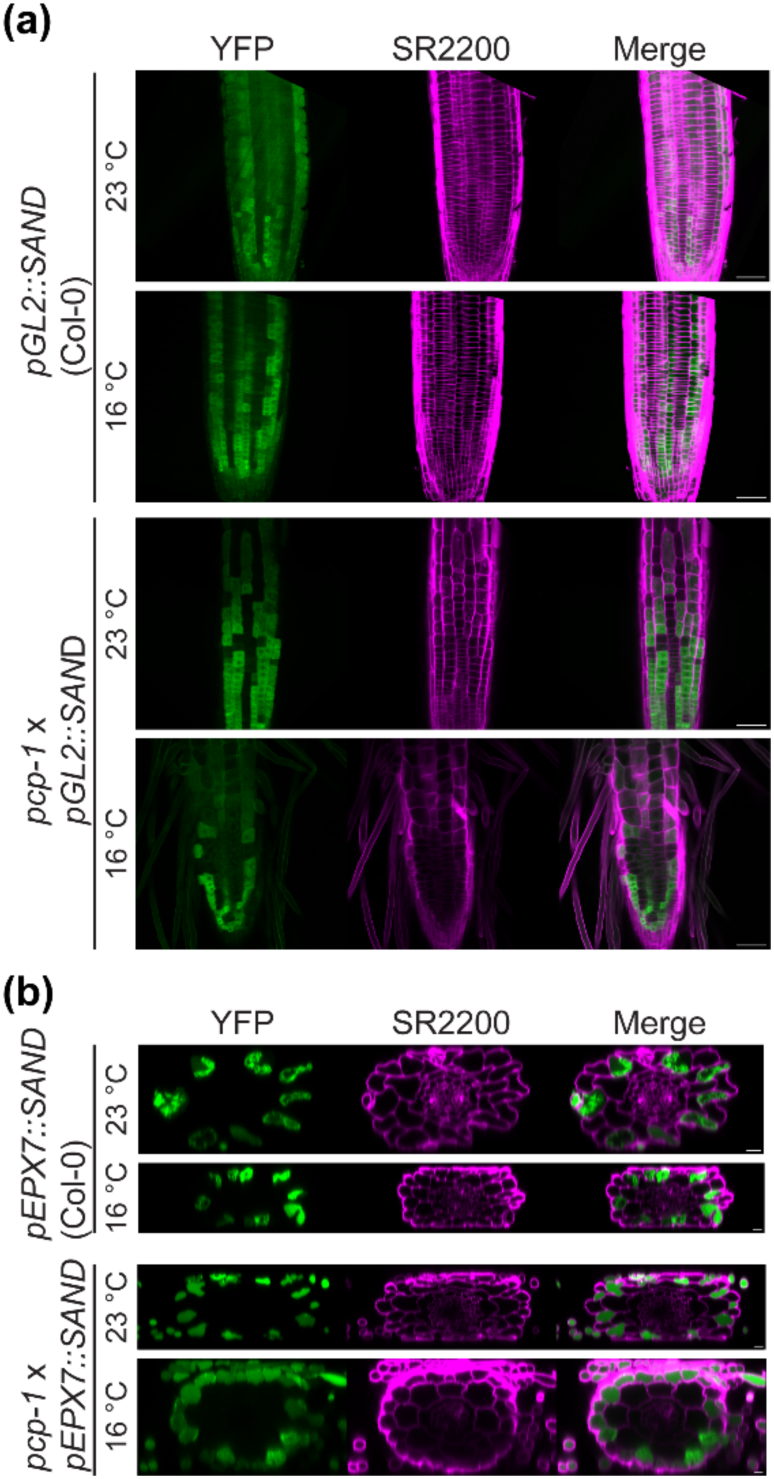
Root hair cell misspecification in *pcp-1* mutants. (a) In Col-0, the atrichoblast marker *pGL2::SAND* is expressed in well-defined cell files. Apparent loss of patterning in *pcp-1* at 16°C. Scale bars: 50 µm. (b) Expression of the trichoblast marker *pEXP7::SAND* is confined to epidermal cells touching two cortical cells in Col-0. *pcp-1* displays ectopic marker expression in epidermal cells touching only one cortical cell. Scale bars: 10 µm.

To investigate the second question, we crossed *pcp-1* with the synthetic auxin response marker *pDR5::SAND* from the SWELLINE collection (Marquès-Bueno *et al*., 2016). In the control, the YFP signal was slightly reduced at 16°C, compared to 23°C, which is in line with previously published data (Zhu *et al*., 2015). Interestingly, we observed that the YFP signal in *pcp-1* increased at 23°C, stretching into the stele, and the signal became wider and stronger at 16°C (Fig. 4**a**). To corroborate these findings, and simultaneously explore whether specific auxin-related pathways were affected in *pcp-1*, we decided to measure auxin content in the roots. Since *pcp-1* roots are very short, we decided to use seedlings grown at 23°C and shifted to 16°C for 2 days. Both previous transcriptomic data (Capovilla *et al*., 2018) and our phenotypic data (this study) supported the idea that 2 days shift to 16°C was sufficient to induce all changes responsible for the *pcp-1* cold-sensitivity. The obtained results showed that IAA concentration was increased in *pcp-1* compared to Col-0 at 23°C and after shift (Fig. 4**b**, Table SS4). Curiously, IAA amounts were also increased in Col-0 after shift, which is contrary to the results obtained from the *pDR5::SAND* marker, and we hypothesize that this may be due to the shift to 16°C. We observed that all measured reversible IAA amino acid or sugar conjugates, which serve as storage/inhibition compounds, were increased, while compounds produced through irreversible oxidative conjugation were increased in *pcp-1* at 23°C, but slightly decreased after shift (Fig. S2**a**). These results prompted us to examine if the elevated auxin content at low temperatures could be responsible for the observed *pcp-1* root phenotype. To this end, we used the known dominant auxin biosynthesis mutant *yuc1D*, which produces elevated amounts of IAA (Zhao *et al*., 2001). While we did observe that *yuc1D* produced an increased amount of root hairs at 16°C, we did not see the same phenotypic aberrations of the RAM as in *pcp-1* (Fig. S2**b**). Overall, these results strengthen the idea that the loss of *PCP* induces changes in auxin homeostasis, which might be compensated at 23°C through the elevated storage/inhibition compounds but may cause more severe phenotypic alterations at 16°C, where oxidative conjugation appears to be attenuated. Furthermore, our results suggest that an increase in IAA at low temperature contributes to the observed root hair phenotype, and thus partially explains the *pcp-1* phenotype, but does not account for all RAM alterations.

**Figure 4.**
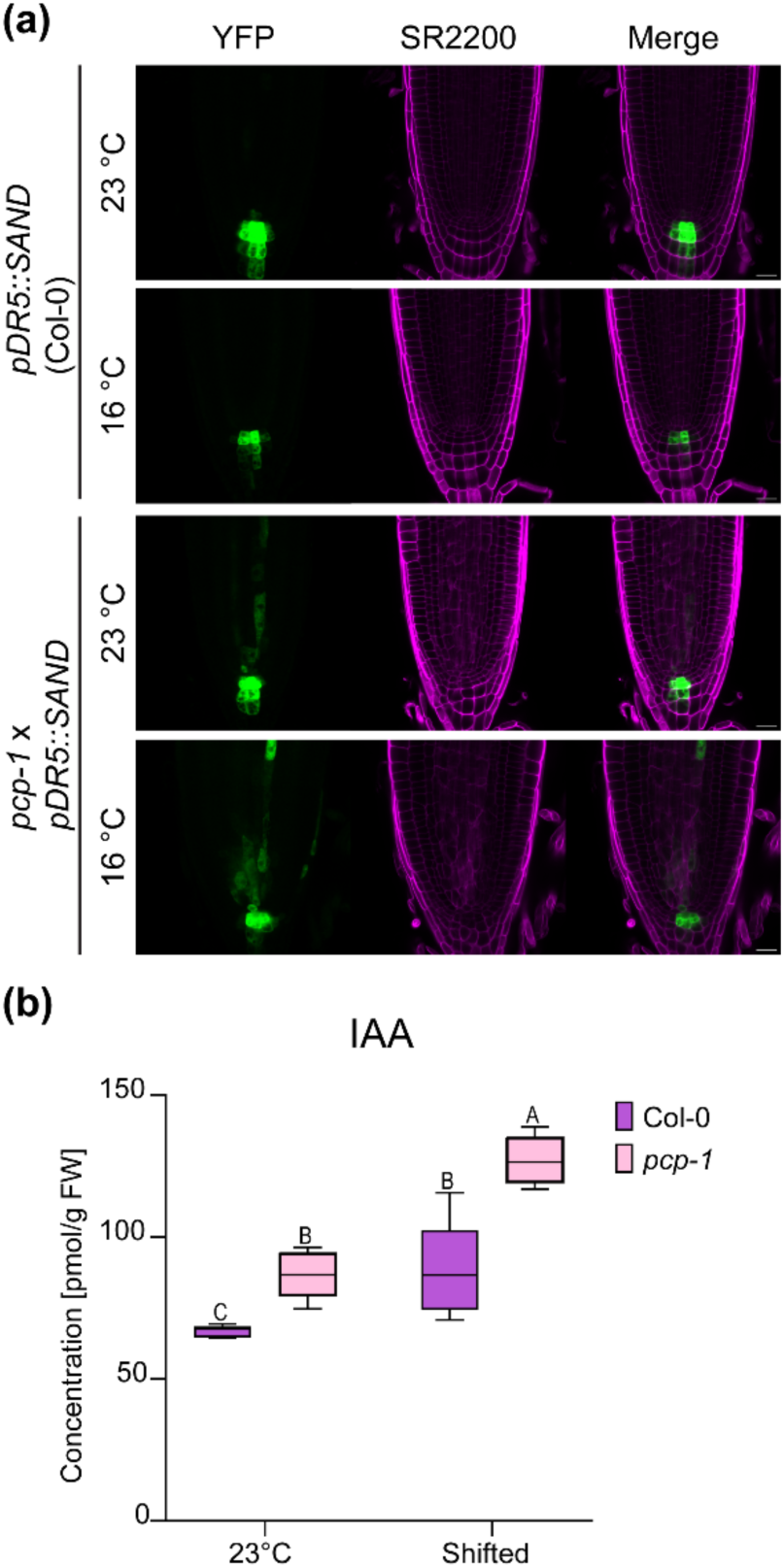
***pcp-1* mutants have higher amounts of IAA in the root meristem.** (a) Transcriptional DR5 auxin response marker shows widened and ectopic activation in *pcp-1* mutants. Scale bars: 20 µm. (b) IAA content is elevated in *pcp-1* seedlings. Box plots are min to max. Statistical test: Two-way ANOVA with posthoc Tukey analysis.

Since our results showed that the endogenous auxin content was elevated and ectopically localized in the stele of *pcp-1*, we decided to explore the possibility of PIN protein mislocalization. To account for auxin flow in the stele, cortex, and columella, we chose to examine PIN1, PIN2, and PIN7 (Ruiz Rosquete *et al*., 2012). At 23°C these PIN proteins localized correctly in both genotypes (Fig. 5**a**). At 16°C all tested PIN proteins localized correctly in Col-0, though a reduced signal strength was observed for PIN7. In *pcp-1* we found that PIN1 and PIN2 localization appeared aberrant at first glance. However, closer examination revealed that PIN1 still localized acropetally, and that the apparent mislocalization was due to the morphological defects of cells in the inner stele, as basal cell walls were not perpendicular to the root-shoot axis. Similarly, PIN2 appeared to be localized basipetally in the epidermis, and cortical PIN2 was found mostly acropetally, but due to the severe defects of cortical cells, the basal cell walls were rather pointing towards the stele. PIN7 signal strength was very low in *pcp-1* at 16°C and undetectable in most seedlings (Fig. 5**b**). When detectable, it was correctly localized in the stele, but nearly undetectable in the columella. Taken together it thus seems possible that aberrant centripetal auxin flow through PIN1 and PIN2, coupled with a lack of PIN7 proteins in the columella could cause an accumulation of auxin in the stele, as observed in *pcp-1*. In summary, these results suggest that PCP regulates patterning processes in the *A. thaliana* root, presumably through the regulation of auxin homeostasis and flow.

**Figure 5.**
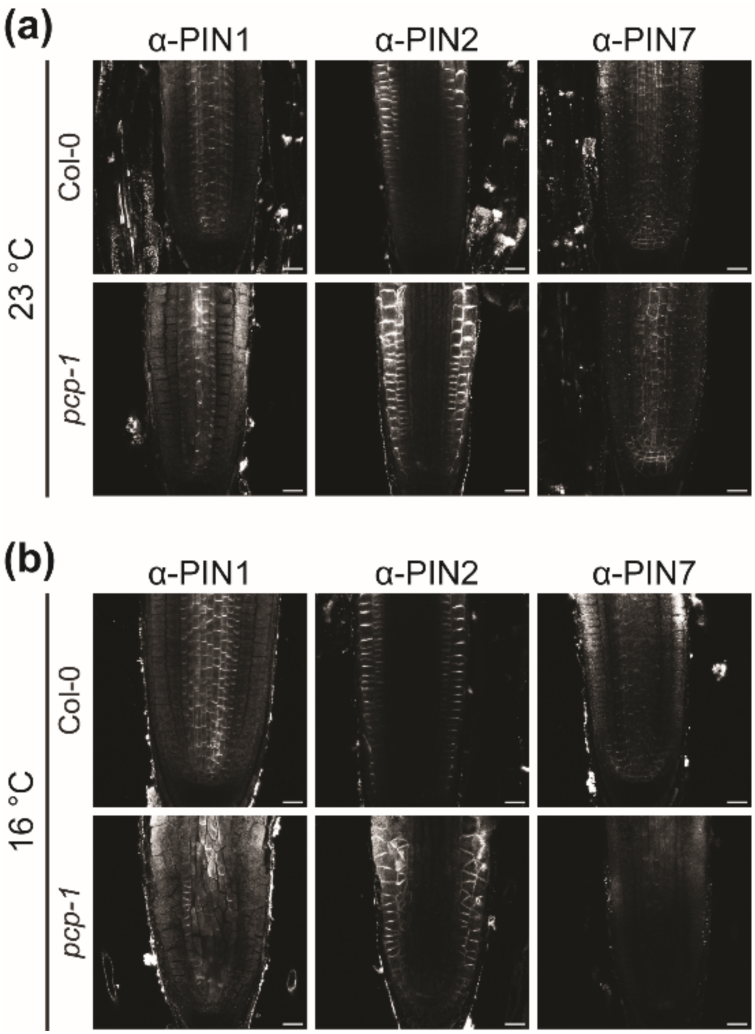
Immunolocalization of PIN proteins reveals that auxin transport in *pcp-1* roots is disturbed. (a) PIN protein localization at 23°C is correct in Col-0 and *pcp-1.* (b) PIN protein localization is correct in Col-0 at 16°C, but PIN7 signal strength appears to be lower than in 23°C. PIN protein localization in *pcp-1* appears aberrant due to cell morphology and only weak PIN7 signal strength is detectable.

### Inhibition of auxin response partially rescues the *pcp-1* RAM phenotype

Our data shows that the loss of *PCP* causes temperature-dependent defects in root development, characterized by ectopic WOX5 expression, root hair misspecification, and an increase in endogenous IAA content. Based on these findings, we decided to treat seedlings with 10 µM auxinole, a known auxin response inhibitor (Hayashi *et al*., 2012), or 100 nM IAA, to mimic the increased auxin content at low temperature. Auxinole treatment had no major effect on the root length of Col-0 or *pcp-1*, while some root waving was observed, which is consistent with previous observations (Hayashi *et al*., 2012). Interestingly, we recorded a decreased root hair density in *pcp-1* grown at 16°C in response to the auxinole treatment (Fig. S3**a**). IAA treatment induced root shortening in both genotypes and temperatures (Fig. S3**b**). We observed, however, that root shortening and increase in root hair density was more pronounced in Col-0 at 16°C, suggesting that elevated IAA and low temperature may have an additive negative effect on root development. We also found that *pcp-1* reacted more sensitively to exogenous IAA at 23°C, which may be a result of either the elevated endogenous IAA content or due to a disruption of other buffering mechanisms. At a microscopic level, we did not observe any discernable differences in RAM architecture in Col-0 in response to auxinole or IAA treatment (Fig. S4**a**). However, our observations revealed that auxinole treatment at 16°C markedly improved RAM architecture in *pcp-1*, while IAA treatment worsened the phenotype, inducing cortical cell swelling. We did not observe any cortical cell swelling at 23°C, implying a temperature-dependent disruption of auxin homeostasis in *pcp-1* mutants (Fig. S4**b**).

Based on these findings, we were curious to see whether WOX5 expression was affected in response to these treatments. We found that auxinole treatment induced a laterally widened area of WOX5 expressing cells in both the control and *pcp-1* at 23°C (Fig. 6**c**, Fig. S4**c**). At 16°C we observed a similar effect, while mutants depicted less ectopic WOX5 expression in the stele (Fig. 6**a**,**c**). Interestingly, while IAA treatment induced no significant changes in GFP signal in the control at either temperature, we observed a significantly higher number of WOX5 positive cells in *pcp-1* crossed with *pWOX5::ER-GFP* at 23°C (Fig. 6**d**, Fig. S4**d**). We observed no significant increase in WOX5 positive cells in *pcp-1* at 16°C (Fig. 6**b**,**d**).

**Figure 6.**
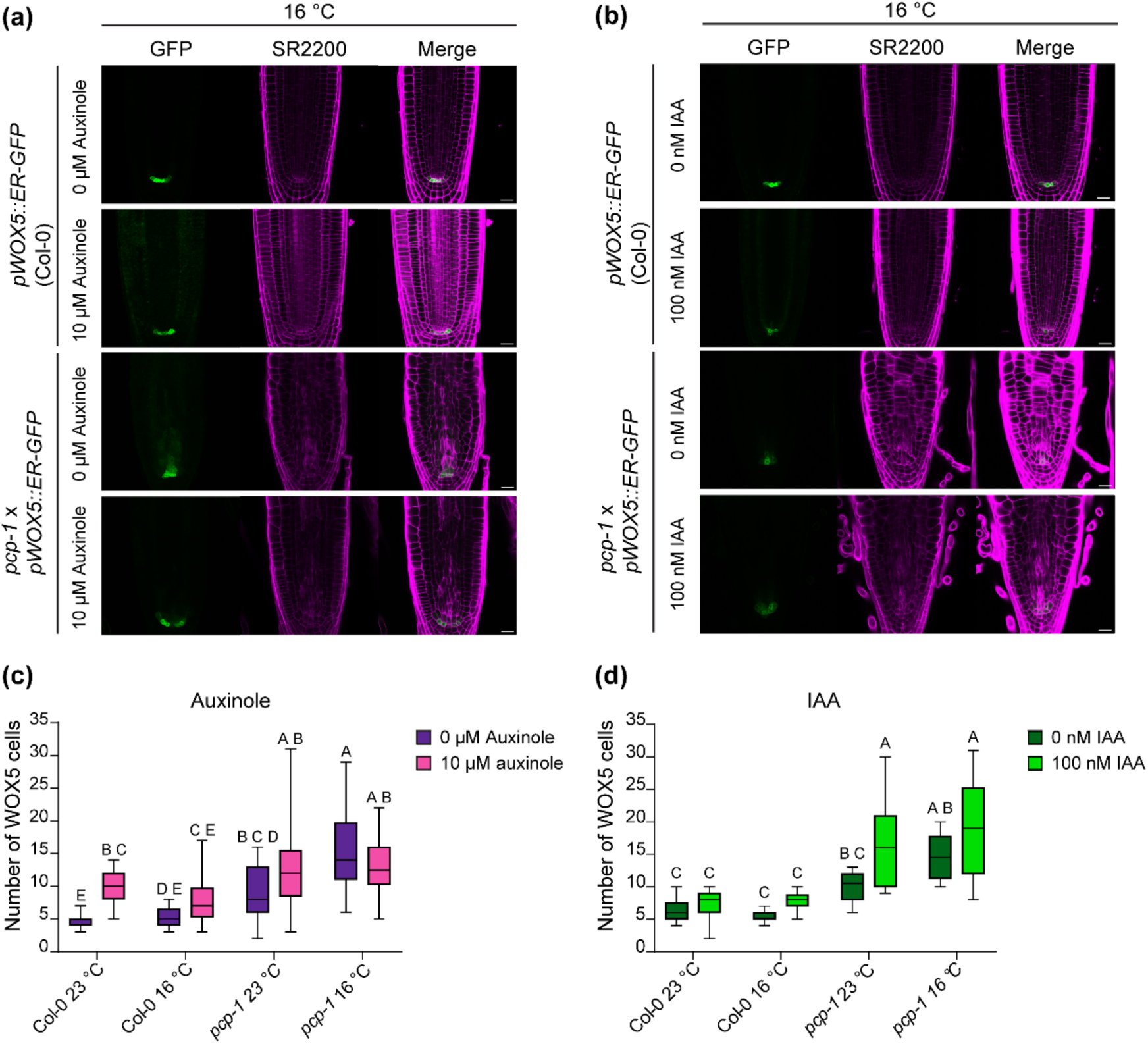
Auxinole treatment partially rescues the *pcp-1* cold-sensitive phenotype, while it is worsened by IAA treatment. (a) Ectopic expression of the *pWOX5::GFP* reporter in *pcp-1* mutants, and root patterning are partially rescued by treatment with 10 µM auxinole. Scale bars: 20 µm. (b) IAA treatment causes cortical cell swelling in *pcp-1*. Scale bars: 20 µm. (c-d) Quantification of *WOX5* expressing cells in auxinole and IAA treated seedlings. Box plots are min to max. Statistical test: Two-way ANOVA with posthoc Tukey analysis.

In conclusion, we found that 10 µM auxinole partially suppressed the *pcp-1* root hair phenotype and improved RAM architecture, while treatment with 100 nM IAA at 16°C induced macroscopic phenotypic similarities between Col-0 and *pcp-1*. Furthermore, auxinole treatment reduced ectopic *WOX5* expression in *pcp-1* at 16°C, while IAA treatment increased it at 23°C, indicating that *pcp-1* mutants may have a defect in auxin response regulation.

### Low-temperature root transcriptomics in response to IAA and auxinole

Auxinole treatment partially rescued *pcp-1* RAM defects at 16°C, while IAA treatment at 16°C induced a macroscopic phenotype in Col-0 which was reminiscent of *pcp-1*. We decided to make use of these treatments to identify genes which are potentially responsible for the developmental defects observed in *pcp-1*. Our mRNA sequencing results showed that only a small number of genes was specifically affected by the treatments, and that most differentially expressed genes were found between genotypes (Fig. 7**a**-**b**). We decided to assess our data for clusters of genes that would fulfill the following criteria: (a) induce expression changes in IAA-treated Col-0, to resemble expression levels in *pcp-1*, and (b) induce expression changes in auxinole-treated *pcp-1*, to resemble Col-0. Using Euclidean distance measurement with the ward.D clustering method implemented in the 3D RNAseq analysis application (Guo *et al*., 2021), we identified one cluster that fulfilled both requirements and showed opposite expression changes in response to the different treatments. Gene ontology (GO) enrichment of this cluster showed that it was, among others, enriched in processes linked to root and root hair development, and response to auxin (Fig. 7**c**-**d**, Table S8). These results were confirmed using DIANE (Cassan *et al*., 2021), which employs the Poisson mixture models clustering, thus modelling the distribution of the underlying data (Gao *et al*., 2023). Using the same criteria as described above, we found that one cluster matched our requirements, and GO enrichment of this cluster gave nearly identical results (Fig. S5**a**-**b**). We decided to also explore the possible network clusters from DIANE and found one cluster, which was also enriched for epidermal cell and root developmental processes (Fig. S5**c**-**d**).

**Figure 7.**
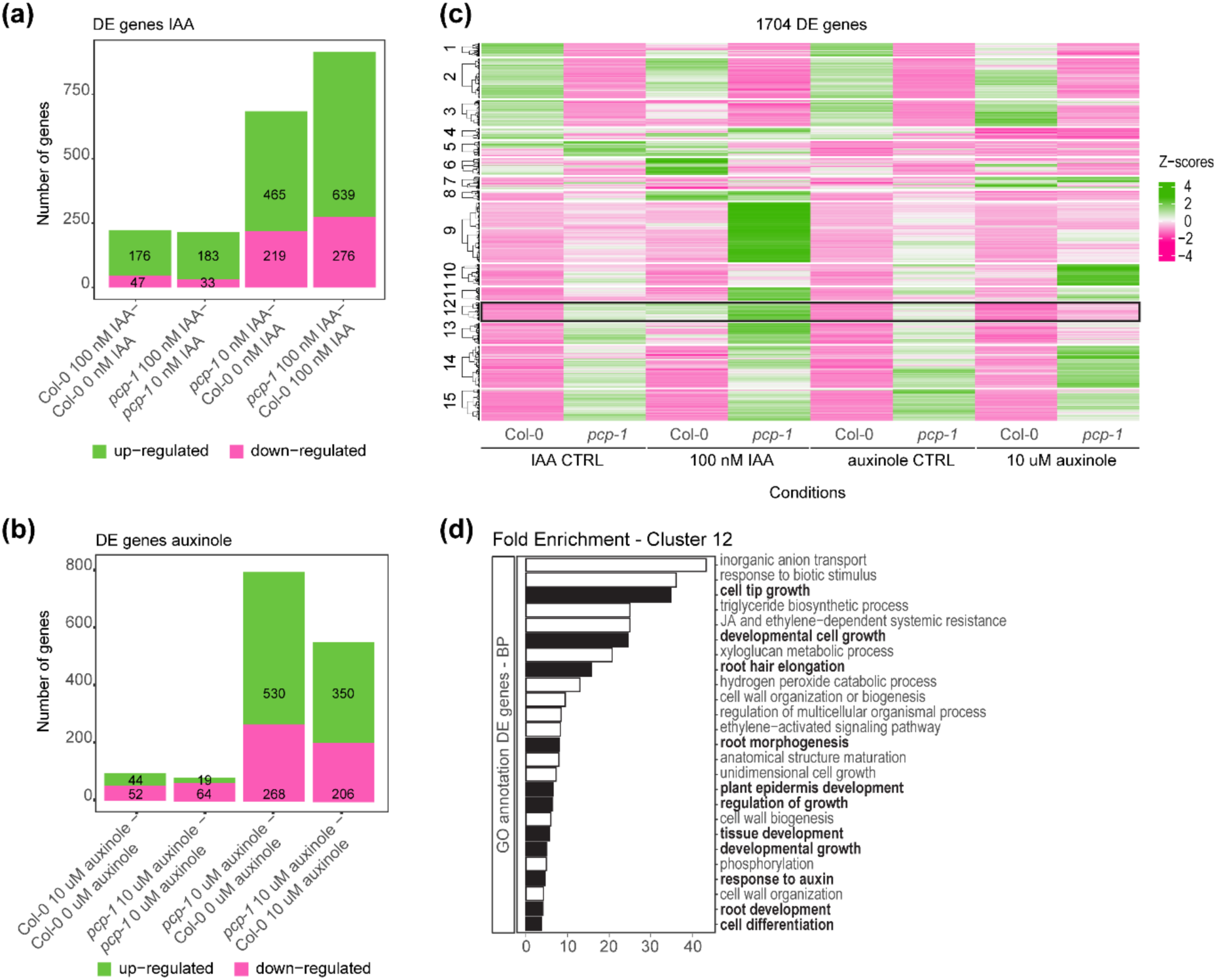
RNA sequencing data reveals gene cluster associated with root morphology. (a-b) Differentially expressed genes are mostly due to genotype. Only a low number of genes is mis-regulated in response to auxinole and/or IAA to treatment. (c) Heatmap of all differentially expressed genes. (d) Genes in cluster 12 of the heatmap are enriched in processes involved in root development, root morphology and hormone response according to gene ontology enrichment (BP: Biological Process, -log10(FDR)) analysis performed in DAVID (Database for Annotation, Visualization and Integrated Discovery). Dark grey bars highlight GO terms of interest.

Finally, we examined whether *WOX5* expression in the RNA sequencing data reflected our previous results. We found that *WOX5* expression was strongly elevated in *pcp-1* and reduced in response to auxinole treatment. The same was found for *WOX7* expression. Finally, *WOX11*, which links auxin signaling and root system architecture (Sheng *et al*., 2017), was elevated in *pcp-1*. While auxinole treatment did not reduce *WOX11* expression, it did revert the expression ratio of some isoforms to that observed in Col-0 (Fig. S6**a**-**c**), suggesting that the splicing of at least some genes, which modulate root architecture, is influenced by the auxin response in *pcp-1*.

## Discussion

Among the multitude of environmental signals, which plants need to adjust their growth and development to, our research focuses on ambient temperature. The relevance of temperature-dependent AS is supported by the findings of Calixto *et al*., 2018, who reported that changes of only 2°C induced measurable differences in AS. Interestingly, this seems to be particularly evident for mutants affected in *PCP*/*SmE1*/*SmEb*, where different degrees of developmental defects were observed in control temperatures ranging from 20°C to 23°C (Capovilla *et al*., 2018; Huertas *et al*., 2019; Wang *et al*., 2022; Willems *et al*., 2023; Hong *et al*., 2023). Notably, Willem *et al*., 2023 observed that the *pcp-1* shoot phenotype disappeared at 26°C, while we observed full suppression of the RAM phenotype at 27°C. Two paralogs of the *SmE* gene can be found in *A. thaliana* (Cao *et al*., 2011), i.e. *PCP* and *PCP-LIKE* (*PCPL*/*SmE2*/*SmEa*/*AT4G30330*), which only differ by two amino acids (Huertas *et al*., 2019). There is some indication that *PCPL* expression could be induced by elevated temperatures (Capovilla *et al*., 2018). Hence, one possible explanation is that the expression of *PCP* and *PCPL* is gradually modulated to provide sufficient SmE protein over a wide temperature range.

The pleiotropic phenotype of *pcp-1* complicates determining the exact mechanisms by which PCP regulates abiotic stress responses. However, using several marker lines, chemical treatments, root hormone profiling and transcriptomic analyses we have aimed to contribute to the understanding of this topic. We found that the loss of *PCP* causes hypersensitivity to low ambient temperatures, marked by a drastic reduction in meristem size, and early differentiation. Our data suggest that cell proliferation is strongly inhibited in *pcp-1* at 16°C, shown by an elevated and ectopic *pWOX5::ER-GFP* expression, as well as reduced expression of mitotic markers. These observations support previous findings, which have shown that *WOX5* overexpression strongly reduces the number of mitotic cells in the proximal meristem (Savina *et al*., 2020). Furthermore, we could show that the observed RAM defects are at least partially linked to elevated endogenous auxin, and the establishment of ectopic auxin maxima in the stele of *pcp-1*. Recent investigations into RAM organization have suggested that WOX5 acts upstream of auxin biosynthetic genes and regulates their expression (Tian *et al*., 2014; Savina *et al*., 2020). The ectopic *WOX5* expression in *pcp-1* could thus contribute to the additional production of auxin and partially explain the ectopic auxin maxima in the vasculature. Finally, our results indicate that elevated auxin and low temperature have an additive negative effect on root development and that the inhibition of auxin response partially suppresses the observed developmental defects in *pcp-1*.

A recent study has shown that WOX5 interacts with PLETHORA (PLT) proteins (Burkart *et al*., 2022). Interestingly, they observed a widening of the WOX5 expression domain in *plt2, plt3* mutants, which is partially reminiscent of our observations in *pcp-1* at 16°C. The study highlighted the interaction of WOX5 and PLT3 in nuclear bodies (NBs), which depends on the prion-like domain (PrD) of PLT3 (Burkart *et al*., 2022). While proteins with PrDs have been shown to be important for plant temperature perception (Jung *et al*., 2020; Legen *et al*., 2024), the (dis-)assembly of NBs is also sensitive to temperature fluctuations (Meyer, 2020). Burkart *et al*., hypothesized that the recruitment of WOX5 to NBs by PLT3 is concentration dependent. In our conditions, *PLT3* expression is mildly elevated in *pcp-1* (Fig. S7), while the inhibition of auxin response reduced *WOX5* (Fig.6, Fig. S6), as well as *PLT3* expression (Fig. S7). This makes it possible that WOX5 may be recruited to NBs in *pcp-1* at 16°C. NBs have various functions and are classified into different groups based on their composition (Muñoz-Díaz & Sáez-Vásquez, 2022), but Burkart *et al*., have shown that PLT3-dependent NBs recruit RNA, which is a strong indication that these are involved in RNA metabolism. Out of the RNA-recruiting NBs, only nuclear speckles are currently known to also recruit transcription factors (Muñoz-Díaz & Sáez-Vásquez, 2022), which thus provides a strong link to the formation of spliceosomal complexes. Furthermore, some nuclear speckle RNA binding proteins have already been identified as regulators of auxin-related developmental pathways and stress response (Bardou *et al*., 2014; Bazin *et al*., 2018). Taken together, the observations from previous studies and the results presented here raise the questions whether (a) WOX5-PLT3 containing NBs are formed in *pcp-1* roots, (b) if their formation is directly auxin responsive, and (c) how, if at all, these influence splicing and/or gene expression regulation in response to temperature.

It appears that other splicing-related proteins also contribute to the regulation of auxin: A recent study has shown that PCP interacts with the Sm-LIKE protein 7 (LSM7). The loss of *LSM7* causes sensitivity to high and low temperatures, which appears to be connected to significant alterations in auxin-related pathways, particularly the production of IAA storage and inactivation compounds (Nardeli *et al*., 2023). While we have not examined the expression of specific *GH3* genes in detail here, it is well known that they are centrally involved in plant stress response (Wojtaczka *et al*., 2022; Luo *et al*., 2023), and it will be interesting to elucidate their connection to AS and temperature response. Another example of the intricate balance between splicing, auxin-related signaling, and RAM maintenance comes from investigations of *MERISTEM-DEFECTIVE* (*MDF*), which is strongly expressed in the QC and encodes for a serine/arginine rich (RS)-domain protein (Casson *et al*., 2009). RS-domain proteins are generally classified as RNA-binding proteins involved in splice site recognition (Sahebi *et al*., 2016; Li *et al*., 2023). Interestingly, MDF was shown to be associated with spliceosomal complexes and located to nuclear speckles (de Luxán-Hernández *et al*., 2022). *mdf-1* displays cell division defects and meristematic cell death, causing strongly impaired root development (de Luxán-Hernández *et al*., 2022), and aberrant radial patterning (Thompson *et al*., 2023). Furthermore, *mdf-1* does not establish a correct auxin maximum, possibly due to strongly decreased levels of PIN2 and PIN4 (Casson *et al*., 2009). This observation was later expanded with transcriptomic analyses, which established that enriched GO categories in *mdf-1* not only relate to, among others, auxin transport, response and signaling (de Luxán-Hernández *et al*., 2022; Thompson *et al*., 2023). It was thus concluded that MDF-mediated splicing may control cell division and development (de Luxán-Hernández *et al*., 2022) as well as promote meristem function and auxin-mediated pathways (Thompson *et al*., 2023). Contrary to these observations, we found that *pcp-1* established ectopic auxin maxima along the stele, particularly at low temperatures. These results were confirmed by our hormone profiling, which showed significantly higher levels of IAA in *pcp-1*. Additionally, our PIN protein localization experiments did not find reduced PIN1 or PIN2 signal strength between Col-0 and *pcp-1*. However, we observed faulty PIN1 and PIN2 localization in *pcp-1* mutants, possibly due to morphological defects in vascular and cortical cells causing a centripetal auxin flow, and reduced PIN7 signal at 16°C. The loss of PIN7-mediated basipetal auxin flow could consequently contribute to auxin accumulation in the root. Taken together, it thus seems that PCP, contrary to MDF, is required to restrict auxin-mediated pathways and suppress meristematic markers. Finally, it is well established that root development is also governed by auxin’s antagonist, cytokinin (Zluhan-Martínez *et al*., 2021), but cytokinin measurements in *pcp-1* were uninformative (Table S5). It has, however, been shown that the loss of the pre-mRNA splicing factor 3 (*RDM16*) causes a reduction in cytokinin levels, while it does not affect auxin (Lv *et al*., 2021). It is, therefore, probable that different splicing factors may regulate alternative hormonal pathways.

The loss of *PCP* also causes early differentiation at low temperatures, highlighted by the early onset of root hairs, and the misspecification of epidermal cells causing an abnormal root hair pattern. Additionally, we observed that *pcp-1* mutants displayed aberrant periclinal and anticlinal cortical cell divisions, which became less severe after treatment with auxinole, and more severe with IAA. Auxin has been shown to have a strong impact on the regulation of cell shape and division through modulation of the cytoskeleton (Vaddepalli *et al*., 2021; García-González & van Gelderen, 2021). In line with these observations, a recent study has shown that activation of cortical auxin response induces anticlinal and periclinal cell divisions, as well as abnormal root hair patterning (Kim *et al*., 2022). While our *pDR5::SAND* auxin response marker showed no activation in the cortex of *pcp-1*, we observed that cortical cell morphology and root hair density was partially rescued in response to auxinole, while root hair density increased in response to IAA in Col-0 at 16°C. The effect of temperature on root hair development is mostly unexplored (Vissenberg *et al*., 2020), thus studying the sum of effects in mutants such as *pcp-1* on root hair development in low temperature could open new research possibilities.

In conclusion, our findings, supported by previous studies, strongly suggest that AS is highly sensitive to minor temperature fluctuations and plays a pivotal role in root meristem maintenance. This is possibly achieved through the sophisticated control of phytohormone homeostasis and meristematic activity.

## Acknowledgments

We thank Dr. Siamsa Doyle and Dr. Ranjan Swarup for their help with the PIN-protein immunolocalization and for providing the required antibodies. We thank Dr. Crisanto Gutiérrez, Dr. Sara Raggi, Dr. Stéphanie Robert, and Dr. Luciano Di Fino for providing the PlaCCI, *yuc1D* and *pWOX5::ER-GFP* seeds, respectively. Research in the Schmid group is funded by the Knut och Alice Wallenberg Stiftelse (KAW 2018.0202) and FORMAS (2023-01077). Research in the Ljung group is funded by the Knut and Alice Wallenberg Foundation (KAW 2016.0352, KAW 2020.0240) and the Swedish Research Council (VR 2021-04938).

Access to instrumentation was provided by the Swedish Metabolomics Centre (https://www.swedishmetabolomicscentre.se/), and the UPSC Microscopy facility (https://www.upsc.se/platforms/microscopy-facility.html). We thank the UPSC Bioinformatics facility (https://www.upsc.se/platforms/upsc-bioinformatics-facility.html) for the bioinformatics support.

## Author contributions

MS, KL: Funding acquisition. MS, NEA, SN: Conceptualization. SN, NEA: Performed experiments and data analysis. MS, NEA: Visualization and writing. JS: Hormone profiling. All authors read and approved the manuscript.

## Data availability

RNA sequencing data for this study have been deposited in the European Nucleotide Archive (ENA) at EMBL-EBI under accession number PRJEBxxxx. Other data supporting this study’s findings, which are not already included in the supplementary material, are available from the corresponding author upon reasonable request.

## Supporting Information – Brief legends

**Fig. S1** Reversion of the *pcp-1* root phenotype at elevated temperatures.

**Fig. S2** Auxin storage and catabolic compounds are elevated in *pcp-1*.

**Fig. S3** Auxinole treatment partially suppresses *pcp-1* root hair phenotype at 16°C, while IAA induces root hair phenotype in Col-0.

**Fig. S4** Auxinole treatment partially rescues root morphology defects in *pcp-1* at 16°C, while IAA appears to induce them.

**Fig. S5** Clustering analysis and network inference of RNA sequencing data.

**Fig. S6** RNA sequencing expression data of meristematic regulators and cell cycle indicators.

**Fig. S7** RNA sequencing expression data *PLT* genes.

**Table S1** *Arabidopsis thaliana* plant material used in this study.

**Table S2** PIN protein immunolocalization: Antibody dilutions

**Table S3** Confocal Microscopy settings.

**Table S4** Statistical Analysis of Auxin Hormone Profiling.

**Table S5** Statistical Analysis of Cytokinin Hormone Profiling.

**Table S6** Statistical Analysis of Root Phenotyping Data.

**Table S7** Statistical Analysis of WOX5 and CycB1;1 cell counts.

**Table S8** RNAseq results 3DApp and DIANE.

**Methods S1** PIN protein immunolocalization in Arabidopsis thaliana roots using the InsituProVSI Robot

## References

Abas L, Benjamins R, Malenica N, Paciorek TT, Wiřniewska J, Moulinier-Anzola JC, Sieberer T, Friml J, Luschnig C. 2006. Intracellular trafficking and proteolysis of the Arabidopsis auxin-efflux facilitator PIN2 are involved in root gravitropism. Nature Cell Biology 2006 8:3 8: 249–256.

Bardou F, Ariel F, Simpson CG, Romero-Barrios N, Laporte P, Balzergue S, Brown JWS, Crespi M. 2014. Long Noncoding RNA Modulates Alternative Splicing Regulators in Arabidopsis. Developmental Cell 30: 166–176.

Bazin J, Romero N, Rigo R, Charon C, Blein T, Ariel F, Crespi M. 2018. Nuclear speckle rna binding proteins remodel alternative splicing and the non-coding arabidopsis transcriptome to regulate a cross-talk between auxin and immune responses. Frontiers in Plant Science 9: 397393.

Bellstaedt J, Trenner J, Lippmann R, Poeschl Y, Zhang X, Friml J, Quint M, Delkera C. 2019. A Mobile Auxin Signal Connects Temperature Sensing in Cotyledons with Growth Responses in Hypocotyls. Plant Physiology 180: 757–766.

Blilou I, Xu J, Wildwater M, Willemsen V, Paponov I, Friml J, Heidstra R, Aida M, Palme K, Scheres B. 2005. The PIN auxin efflux facilitator network controls growth and patterning in Arabidopsis roots. Nature 433: 39–44.

Bolger AM, Lohse M, Usadel B. 2014. Trimmomatic: a flexible trimmer for Illumina sequence data. Bioinformatics 30: 2114–2120.

Burkart RC, Strotmann VI, Kirschner GK, Akinci A, Czempik L, Dolata A, Maizel A, Weidtkamp-Peters S, Stahl Y. 2022. PLETHORA-WOX5 interaction and subnuclear localization control Arabidopsis root stem cell maintenance. EMBO reports 23.

Calixto CPG, Guo W, James AB, Tzioutziou NA, Entizne JC, Panter PE, Knight H, Nimmo HG, Zhang R, Brown JWS. 2018. Rapid and Dynamic Alternative Splicing Impacts the Arabidopsis Cold Response Transcriptome. The Plant Cell 30: 1424–1444.

Cao J, Shi F, Liu X, Jia J, Zeng J, Huang G. 2011. Genome-Wide Identification and Evolutionary Analysis of Arabidopsis Sm Genes Family. Journal of Biomolecular Structure and Dynamics 28: 535–544.

Capovilla G, Delhomme N, Collani S, Shutava I, Bezrukov I, Symeonidi E, de Francisco Amorim M, Laubinger S, Schmid M. 2018. PORCUPINE regulates development in response to temperature through alternative splicing. Nature Plants 4: 534–539.

Casanova-Sáez R, Mateo-Bonmatí E, Ljung K. 2021. Auxin Metabolism in Plants. Cold Spring Harbor Perspectives in Biology 13: a039867.

Casanova-Sáez R, Mateo-Bonmatí E, Šimura J, Pěnčík A, Novák O, Staswick P, Ljung K. 2022. Inactivation of the entire Arabidopsis group II GH3s confers tolerance to salinity and water deficit. New Phytologist 235: 263–275.

Cassan O, Lèbre S, Martin A. 2021. Inferring and analyzing gene regulatory networks from multi-factorial expression data: a complete and interactive suite. BMC Genomics 22: 1–15.

Casson SA, Topping JF, Lindsey K. 2009. MERISTEM-DEFECTIVE, an RS domain protein, is required for the correct meristem patterning and function in Arabidopsis. The Plant Journal 57: 857–869.

Danve C, Castroverde M, Dina D, Van Zanten M. 2021. Temperature regulation of plant hormone signaling during stress and development. Journal of Experimental Botany 72: 7436– 7458.

Desvoyes B, Arana-Echarri A, Barea MD, Gutierrez C. 2020. A comprehensive fluorescent sensor for spatiotemporal cell cycle analysis in Arabidopsis. Nature Plants 2020 6:11 6: 1330–1334.

Dikaya V, El Arbi N, Rojas-Murcia N, Muniz Nardeli S, Goretti D, Schmid M. 2021. Insights into the role of alternative splicing in plant temperature response. Journal of Experimental Botany 72: 7384–7403.

Doyle SM, Haegera A, Vain T, Rigala A, Viotti C, Łangowskaa M, Maa Q, Friml J, Raikhel N V., Hickse GR, et al. 2015. An early secretory pathway mediated by gnom-like 1 and gnom is essential for basal polarity establishment in Arabidopsis thaliana. Proceedings of the National Academy of Sciences of the United States of America 112: E806–815.

Doyle SM, Rigal A, Grones P, Karady M, Barange DK, Majda M, Pařízková B, Karampelias M, Zwiewka M, Pěnčík A, et al. 2019. A role for the auxin precursor anthranilic acid in root gravitropism via regulation of PIN-FORMED protein polarity and relocalisation in Arabidopsis. New Phytologist 223: 1420–1432.

Forzani C, Aichinger E, Sornay E, Willemsen V, Laux T, Dewitte W, Murray JAH. 2014. WOX5 Suppresses CYCLIN D Activity to Establish Quiescence at the Center of the Root Stem Cell Niche. Current Biology 24: 1939–1944.

Gao CX, Dwyer D, Zhu Y, Smith CL, Du L, Filia KM, Bayer J, Menssink JM, Wang T, Bergmeir C, et al. 2023. An overview of clustering methods with guidelines for application in mental health research. Psychiatry Research 327: 115265.

García-González J, van Gelderen K. 2021. Bundling up the Role of the Actin Cytoskeleton in Primary Root Growth. Frontiers in Plant Science 12: 777119.

González-García MP, Conesa CM, Lozano-Enguita A, Baca-González V, Simancas B, Navarro-Neila S, Sánchez-Bermúdez M, Salas-González I, Caro E, Castrillo G, et al. 2023. Temperature changes in the root ecosystem affect plant functionality. Plant Communications 4.

Greb T, Lohmann JU. 2016. Plant Stem Cells. Current Biology 26: R816–R821.

Guo X, Liu D, Chong K. 2018. Cold signaling in plants: Insights into mechanisms and regulation. Journal of Integrative Plant Biology 60: 745–756.

Guo W, Tzioutziou NA, Stephen G, Milne I, Calixto CPG, Waugh R, Brown JWS, Zhang R. 2021. 3D RNA-seq: a powerful and flexible tool for rapid and accurate differential expression and alternative splicing analysis of RNA-seq data for biologists. RNA Biology 18: 1574–1587.

Hannah MA, Wiese D, Freund S, Fiehn O, Heyer AG, Hincha DK. 2006. Natural Genetic Variation of Freezing Tolerance in Arabidopsis. Plant Physiology 142: 98–112.

Hayashi K-I, Arai K, Aoi Y, Tanaka Y, Hira H, Guo R, Hu Y, Ge C, Zhao Y, Kasahara H, et al. 2021. The main oxidative inactivation pathway of the plant hormone auxin. Nature communications 12: 6752.

Hayashi KI, Neve J, Hirose M, Kuboki A, Shimada Y, Kepinski S, Nozaki H. 2012. Rational design of an auxin antagonist of the SCF TIR1 auxin receptor complex. ACS Chemical Biology 7: 590–598.

Hernandez JS, Dziubek D, Schröder L, Seydel C, Kitashova A, Brodsky V, Nägele T. 2023. Natural variation of temperature acclimation of Arabidopsis thaliana. Physiologia Plantarum 175: e14106.

Hong Y, Gao Y, Pang J, Shi H, Li T, Meng H, Kong D, Chen Y, Zhu J, Wang Z. 2023. The Sm core protein SmEb regulates salt stress responses through maintaining proper splicing of *RCD1* pre-mRNA in *Arabidopsis*. Journal of Integrative Plant Biology 65: 1383– 1393.

Huertas R, Catalá R, Jiménez-Gómez JM, Castellano MM, Crevillén P, Piñeiro M, Jarillo JA, Salinas J. 2019. Arabidopsis SME1 regulates plant development and response to abiotic stress by determining spliceosome activity Specificity. Plant Cell 31: 537–554.

Jung JH, Barbosa AD, Hutin S, Kumita JR, Gao M, Derwort D, Silva CS, Lai X, Pierre E, Geng F, et al. 2020. A prion-like domain in ELF3 functions as a thermosensor in Arabidopsis. Nature 2020 585:7824 585: 256–260.

Jung JKH, McCouch S. 2013. Getting to the roots of it: Genetic and hormonal control of root architecture. Frontiers in plant science 4: 186.

Kim H, Jang J, Seomun S, Yoon Y, Jang G. 2022. Division of cortical cells is regulated by auxin in Arabidopsis roots. Frontiers in Plant Science 13: 953225.

Kondo Y, Tamaki T, Fukuda H. 2014. Regulation of xylem cell fate. Frontiers in Plant Science 5: 98650.

Kopylova E, Noé L, Touzet H. 2012. SortMeRNA: fast and accurate filtering of ribosomal RNAs in metatranscriptomic data. Bioinformatics 28: 3211–3217.

Korasick DA, Enders TA, Strader LC. 2013. Auxin biosynthesis and storage forms. Journal of Experimental Botany 64: 2541–2555.

Kuriakose S V., Silvester N. 2016. Genetic and molecular mechanisms of post-embryonic root radial patterning. Indian Journal of Plant Physiology 21: 457–476.

Kurihara D, Mizuta Y, Sato Y, Higashiyama T. 2015. ClearSee: A rapid optical clearing reagent for whole-plant fluorescence imaging. Development (Cambridge) 142: 4168–4179.

Laloum T, Martín G, Duque P. 2018. Alternative Splicing Control of Abiotic Stress Responses. Trends in Plant Science 23: 140–150.

Lamers J, Der Meer T Van, Testerink C. 2020. How plants sense and respond to stressful environments. Plant Physiology 182: 1624–1635.

Lasok H, Nziengui H, Kochersperger P, Ditengou FA. 2023. Arabidopsis Root Development Regulation by the Endogenous Folate Precursor, Para-Aminobenzoic Acid, via Modulation of the Root Cell Cycle. Plants 12.

Legen J, Lenzen B, Kachariya N, Feltgen S, Gao Y, Mergenthal S, Weber W, Klotzsch E, Zoschke R, Sattler M, et al. 2024. A prion-like domain is required for phase separation and chloroplast RNA processing during cold acclimation in Arabidopsis. The Plant Cell koae145.

Li D, Yu W, Lai M. 2023. Towards understandings of serine/arginine-rich splicing factors. Acta Pharmaceutica Sinica B 13: 3181–3207.

Ludwig-Müller J. 2011. Auxin conjugates: their role for plant development and in the evolution of land plants. Journal of Experimental Botany 62: 1757–1773.

Luo P, Li T-T, Shi W-M, Ma Q, Di D-W. 2023. The Roles of GRETCHEN HAGEN3 (GH3)-Dependent Auxin Conjugation in the Regulation of Plant Development and Stress Adaptation. Plants 12: 4111.

de Luxán-Hernández C, Lohmann J, Tranque E, Chumova J, Binarova P, Salinas J, Weingartner M. 2022. MDF is a conserved splicing factor and modulates cell division and stress response in Arabidopsis. Life Science Alliance 6: e202201507.

Lv B, Hu K, Tian T, Wei K, Zhang F, Jia Y, Tian H, Ding Z. 2021. The pre-mRNA splicing factor RDM16 regulates root stem cell maintenance in Arabidopsis. Journal of Integrative Plant Biology 63: 662–678.

Marquès-Bueno MM, Morao AK, Cayrel A, Platre MP, Barberon M, Caillieux E, Colot V, Jaillais Y, Roudier F, Vert G. 2016. A versatile Multisite Gateway-compatible promoter and transgenic line collection for cell type-specific functional genomics in Arabidopsis. The Plant Journal 85: 320–333.

Matera AG, Wang Z. 2014. A day in the life of the spliceosome. Nature Reviews Molecular Cell Biology 15: 108–121.

Meyer HM. 2020. In search of function: nuclear bodies and their possible roles as plant environmental sensors. Current Opinion in Plant Biology 58: 33–40.

Motte H, Vanneste S, Beeckman T. 2019. Molecular and Environmental Regulation of Root Development. Annual Review of Plant Biology 70: 465–488.

Muñoz-Díaz E, Sáez-Vásquez J. 2022. Nuclear dynamics: Formation of bodies and trafficking in plant nuclei. Frontiers in Plant Science 13: 984163.

Musielak T, Bürgel P, Kolb M, Bayer M. 2016. Use of SCRI Renaissance 2200 (SR2200) as a Versatile Dye for Imaging of Developing Embryos, Whole Ovules, Pollen Tubes and Roots. BIO-PROTOCOL 6.

Nardeli SM, Zacharaki V, Rojas-Murcia N, Collani S, Wang K, Bayer M, Schmid M, Goretti D. 2023. Temperature-dependent regulation of Arabidopsis thaliana growth and development by LSM7. bioRxiv: 2023.03.28.534379.

Neumann A, Meinke S, Goldammer G, Strauch M, Schubert D, Timmermann B, Heyd F, Preußner M. 2020. Alternative splicing coupled mRNA decay shapes the temperature-dependent transcriptome. EMBO reports 21.

Olatunji D, Geelen D, Verstraeten I. 2017. Control of Endogenous Auxin Levels in Plant Root Development. International Journal of Molecular Sciences 2017, Vol. 18, Page 2587 18: 2587.

Otero S, Desvoyes B, Peiró R, Gutierrez C. 2016. Histone H3 Dynamics Reveal Domains with Distinct Proliferation Potential in the Arabidopsis Root. The Plant Cell 28: 1361.

Patro R, Duggal G, Love MI, Irizarry RA, Kingsford C. 2017. Salmon provides fast and bias-aware quantification of transcript expression. Nature Methods 2017 14:4 14: 417–419.

Pavelescu I, Vilarrasa-Blasi J, Planas-Riverola A, González-García M, Caño-Delgado AI, Ibañes M. 2018. A Sizer model for cell differentiation in Arabidopsis thaliana root growth. Molecular Systems Biology 14: 7687.

Pênčík A, Casanova-Sáez R, Pilařová V, Žukauskaite A, Pinto R, Micol JL, Ljung K, Novák O. 2018. Ultra-rapid auxin metabolite profiling for high-throughput mutant screening in Arabidopsis. Journal of Experimental Botany 69: 2569–2579.

Penfield S, Warner S, Wilkinson L. 2021. Molecular responses to chilling in a warming climate and their impacts on plant reproductive development and yield. Journal of Experimental Botany 72: 7374–7383.

Praat M, De Smet I, Van Zanten M. 2021. Protein kinase and phosphatase control of plant temperature responses. Journal of Experimental Botany 72: 7459–7473.

Quint M, Delker C, Franklin KA, Wigge PA, Halliday KJ, van Zanten M. 2016. Molecular and genetic control of plant thermomorphogenesis. Nature plants 2: 15190.

Reddy ASN, Marquez Y, Kalyna M, Barta A. 2013. Complexity of the Alternative Splicing Landscape in Plants. The Plant Cell 25: 3657–3683.

Ruiz Rosquete M, Barbez E, Rgen Kleine-Vehn J. 2012. Cellular Auxin Homeostasis: Gatekeeping Is Housekeeping.

Sahebi M, Hanafi MM, van Wijnen AJ, Azizi P, Abiri R, Ashkani S, Taheri S. 2016. Towards understanding pre-mRNA splicing mechanisms and the role of SR proteins. Gene 587: 107–119.

Salazar-Henao JE, Vélez-Bermúdez IC, Schmidt W. 2016. The regulation and plasticity of root hair patterning and morphogenesis. Development 143: 1848–1858.

Sarkar AK, Luijten M, Miyashima S, Lenhard M, Hashimoto T, Nakajima K, Scheres B, Heidstra R, Laux T. 2007. Conserved factors regulate signalling in Arabidopsis thaliana shoot and root stem cell organizers. Nature 2007 446:7137 446: 811–814.

Sauer M, Paciorek T, Benkovó E, Friml J. 2006. Immunocytochemical techniques for whole-mount in situ protein localization in plants. Nature Protocols 2006 1:1 1: 98–103.

Savina MS, Pasternak T, Omelyanchuk NA, Novikova DD, Palme K, Mironova V V., Lavrekha V V. 2020. Cell Dynamics in WOX5-Overexpressing Root Tips: The Impact of Local Auxin Biosynthesis. Frontiers in Plant Science 11: 560169.

Sheng L, Hu X, Du Y, Zhang G, Huang H, Scheres B, Xu L. 2017. Non-canonical WOX11-mediated root branching contributes to plasticity in arabidopsis root system architecture. Development (Cambridge) 144: 3126–3133.

Šimura J, Antoniadi I, Široká J, Tarkowská D, Strnad M, Ljung K, Novák O. 2018. Plant Hormonomics: Multiple Phytohormone Profiling by Targeted Metabolomics. Plant Physiology 177: 476–489.

De Smet I, Quint M, van Zanten M. 2021. High and low temperature signalling and response. Journal of Experimental Botany 72: 7339–7344.

Staiger D, Brown JWS. 2013. Alternative Splicing at the Intersection of Biological Timing, Development, and Stress Responses. The Plant Cell 25: 3640–3656.

Thompson HL, Shen W, Matus R, Kakkar M, Jones C, Dolan D, Grellscheid S, Yang X, Zhang N, Mozaffari-Jovin S, et al. 2023. MERISTEM-DEFECTIVE regulates the balance between stemness and differentiation in the root meristem through RNA splicing control. Development (Cambridge, England) 150.

Tian H, Wabnik K, Niu T, Li H, Yu Q, Pollmann S, Vanneste S, Govaerts W, Rolčík J, Geisler M, et al. 2014. WOX5–IAA17 Feedback Circuit-Mediated Cellular Auxin Response Is Crucial for the Patterning of Root Stem Cell Niches in Arabidopsis. Molecular Plant 7: 277–289.

Tofanelli R, Vijayan A, Scholz S, Schneitz K. 2019. Protocol for rapid clearing and staining of fixed Arabidopsis ovules for improved imaging by confocal laser scanning microscopy. Plant Methods 15: 1–13.

Vaddepalli P, de Zeeuw T, Strauss S, Bürstenbinder K, Liao CY, Ramalho JJ, Smith RS, Weijers D. 2021. Auxin-dependent control of cytoskeleton and cell shape regulates division orientation in the Arabidopsis embryo. Current Biology 31: 4946–4955.e4.

Verbelen JP, De Cnodder T, Le J, Vissenberg K, Baluška F. 2006. The Root Apex of Arabidopsis thaliana Consists of Four Distinct Zones of Growth Activities: Meristematic Zone, Transition Zone, Fast Elongation Zone and Growth Terminating Zone. Plant Signaling & Behavior 1: 296.

Vissenberg K, Claeijs N, Balcerowicz D, Schoenaers S. 2020. Hormonal regulation of root hair growth and responses to the environment in Arabidopsis. Journal of Experimental Botany 71: 2412–2427.

Wang Z, Hong Y, Yao J, Huang H, Qian B, Liu X, Chen Y, Pang J, Zhan X, Zhu J, et al. 2022. Modulation of plant development and chilling stress responses by alternative splicing events under control of the spliceosome protein SmEb in *Arabidopsis*. *Plant*, Cell & Environment 45: 2762–2779.

Willems P, Van Ruyskensvelde V, Maruta T, Pottie R, Fernández-Fernández ÁD, Pauwels J, Hannah MA, Gevaert K, Van Breusegem F, Van der Kelen K. 2023. Mutation of Arabidopsis SME1 and Sm core assembly improves oxidative stress resilience. Free Radical Biology and Medicine 200: 117–129.

Wojtaczka P, Ciarkowska A, Starzynska E, Ostrowski M. 2022. The GH3 amidosynthetases family and their role in metabolic crosstalk modulation of plant signaling compounds. Phytochemistry 194: 113039.

Zhang R, Calixto CPG, Marquez Y, Venhuizen P, Tzioutziou NA, Guo W, Spensley M, Entizne JC, Lewandowska D, Have S Ten, et al. 2017. A high quality Arabidopsis transcriptome for accurate transcript-level analysis of alternative splicing. Nucleic Acids Research 45: 5061.

Zhao Y, Christensen SK, Fankhauser C, Cashman JR, Cohen JD, Weigel D, Chory J. 2001. A role for flavin monooxygenase-like enzymes in auxin biosynthesis. Science 291: 306–309.

Zhu T, van Zanten M, De Smet I. 2022. Wandering between hot and cold: temperature dose-dependent responses. Trends in Plant Science 27: 1124–1133.

Zhu J, Zhang KX, Wang WS, Gong W, Liu WC, Chen HG, Xu HH, Lu YT. 2015. Low Temperature Inhibits Root Growth by Reducing Auxin Accumulation via ARR1/12. Plant and Cell Physiology 56: 727–736.

Zluhan-Martínez E, López-Ruíz BA, García-Gómez ML, García-Ponce B, de la Paz Sánchez M, Álvarez-Buylla ER, Garay-Arroyo A. 2021. Integrative Roles of Phytohormones on Cell Proliferation, Elongation and Differentiation in the Arabidopsis thaliana Primary Root. Frontiers in Plant Science 12: 659155.

