## Supplementary Figures 1-7, Tables 1-3 for "The *Arabidopsis* splicing factor PORCUPINE/SmE1 orchestrates temperature-dependent root development via auxin homeostasis maintenance"

New Phytologist

**_New Phytologist_ Supporting Information**

Article acceptance date: Click here to enter a date.

The following Supporting Information is available for this article:

**Fig. S1** Reversion of the *pcp-1* root phenotype at elevated temperatures.
**Fig. S2** Auxin storage and catabolic compounds are elevated in *pcp-1*.
**Fig. S3** Auxinole treatment partially suppresses *pcp-1* root hair phenotype at 16°C, while IAA induces root hair phenotype in Col-0.
**Fig. S4** Auxinole treatment partially rescues root morphology defects in *pcp-1* at 16°C, while IAA appears to induce them.
**Fig. S5** Clustering analysis and network inference of RNA sequencing data.
**Fig. S6** RNA sequencing expression data of meristematic regulators and cell cycle indicators.
**Fig. S7** RNA sequencing expression data *PLT* genes.
**Table S1** *Arabidopsis thaliana* plant material used in this study.
**Table S2** PIN protein immunolocalization: Antibody dilutions
**Table S3** Confocal Microscopy settings.
**Table S4** Statistical Analysis of Auxin Hormone Profiling.
**Table S5** Statistical Analysis of Cytokinin Hormone Profiling.
**Table S6** Statistical Analysis of Root Phenotyping Data.
**Table S7** Statistical Analysis of WOX5 and CycB1;1 cell counts.
**Table S8** RNAseq results 3DApp and DIANE.
**Methods S1** PIN protein immunolocalization in *Arabidopsis thaliana* roots using the InsituProVSI Robot

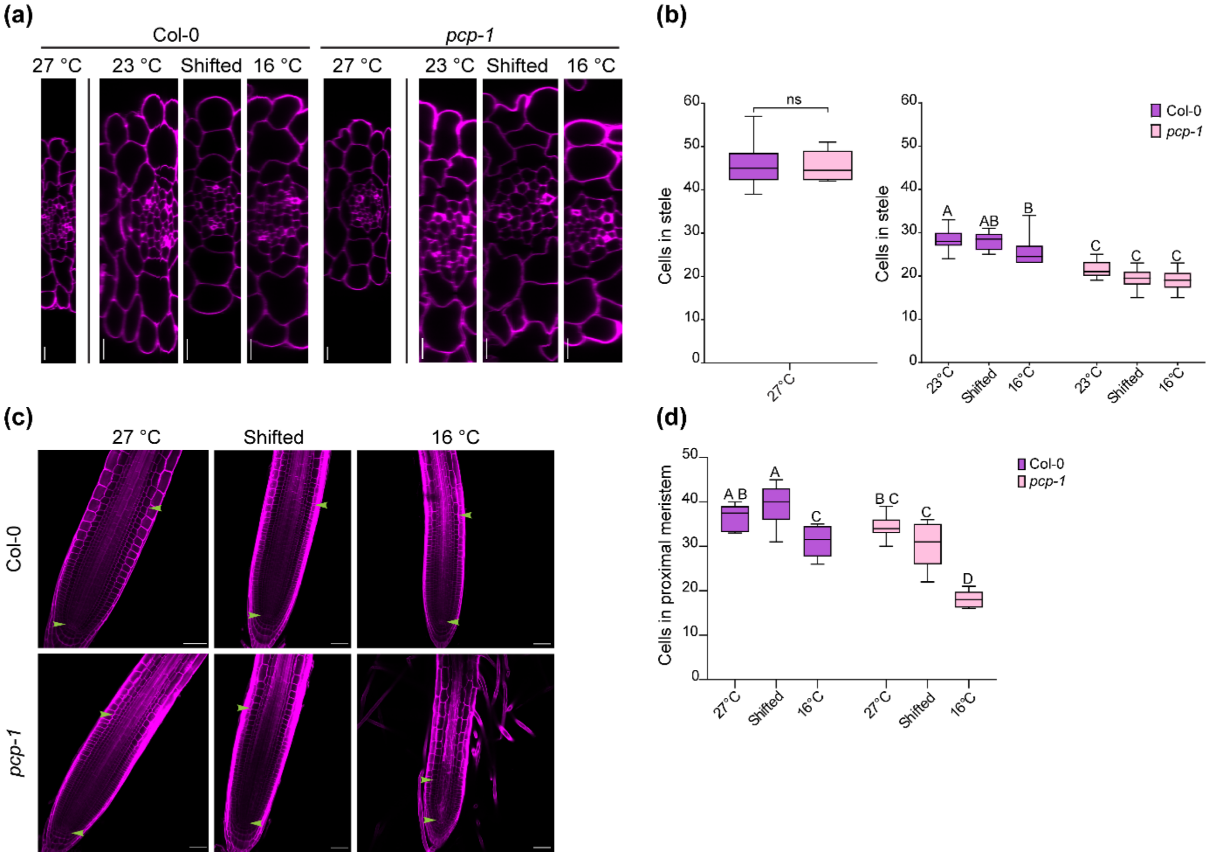


**Fig. S1 Reversion of the *pcp-1* root phenotype at elevated temperatures.**

(a-b) At 23°C and 16°C *pcp-1* has significantly lower number of cells in the stele compared to Col-0. At 27°C no difference in stele cell number is observed. Scale bars: 10 µm. Box plots are min to max. Statistical test: Stele cell counts 27°C: unpaired, two-tailed t-test; Stele cell counts 23°C, shift and 16°C: Two-way ANOVA with posthoc Tukey analysis. (c-d) At 27°C the number of cells in the proximal meristem is not significantly different between Col-0 and *pcp-1*. Scale bars: 50 µm. Box plots are min to max. Statistical test: Two-way ANOVA with posthoc Tukey analysis.

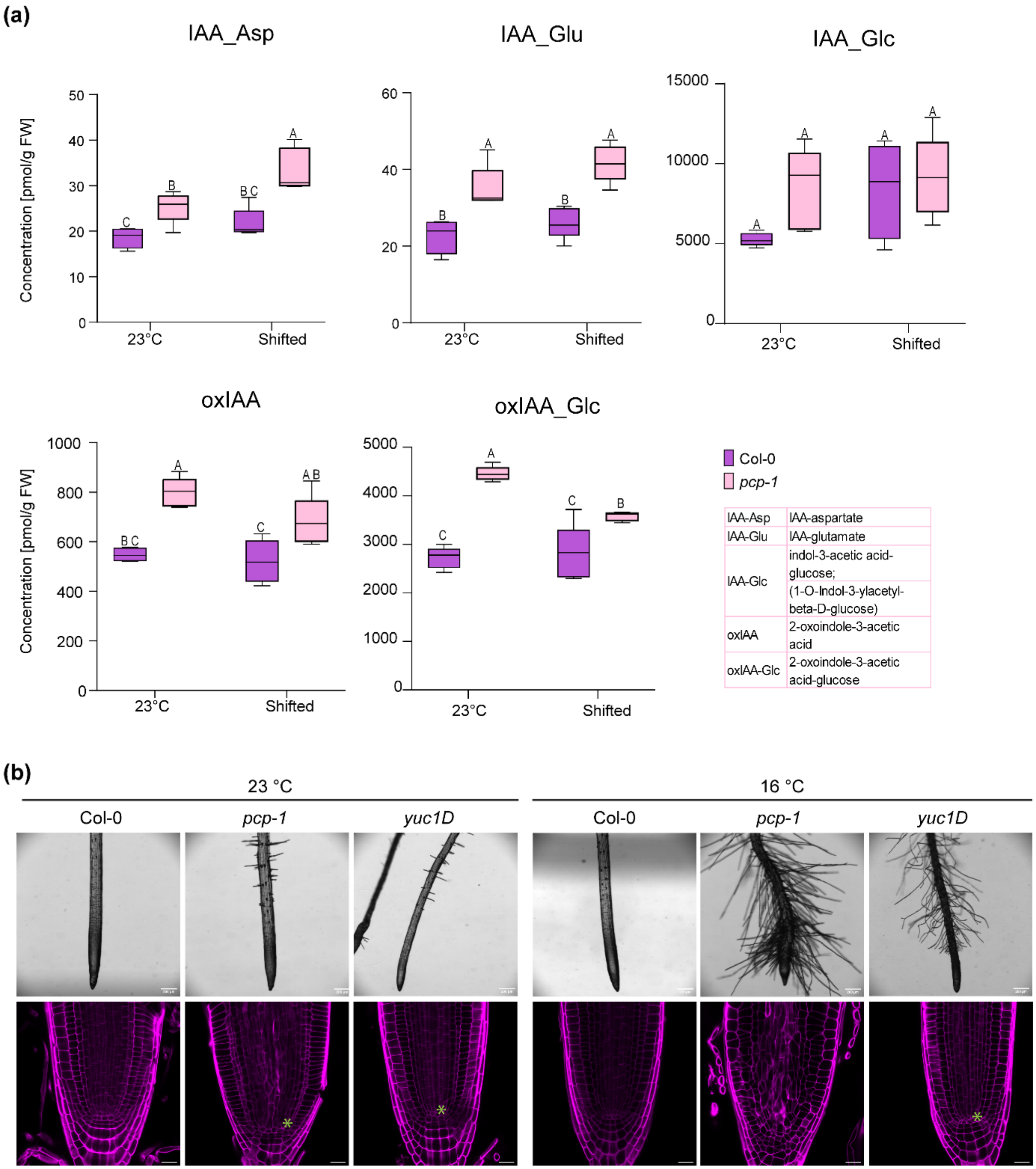


**Fig. S2 Auxin storage and catabolic compounds are elevated in *pcp-1*.**
(a) All measured auxin conjugates and catabolites are elevated in *pcp-1*. Box plots are min to max. Statistical test: Two-way ANOVA with posthoc Tukey analysis. (b) The dominant auxin biosynthesis mutant *yuc1D* produces more root hairs at 16°C. No similarity to the *pcp-1* RAM defects could be observed in *yuc1D*. Scale bars top: 200 µm. Scale bars bottom: 20 µm.

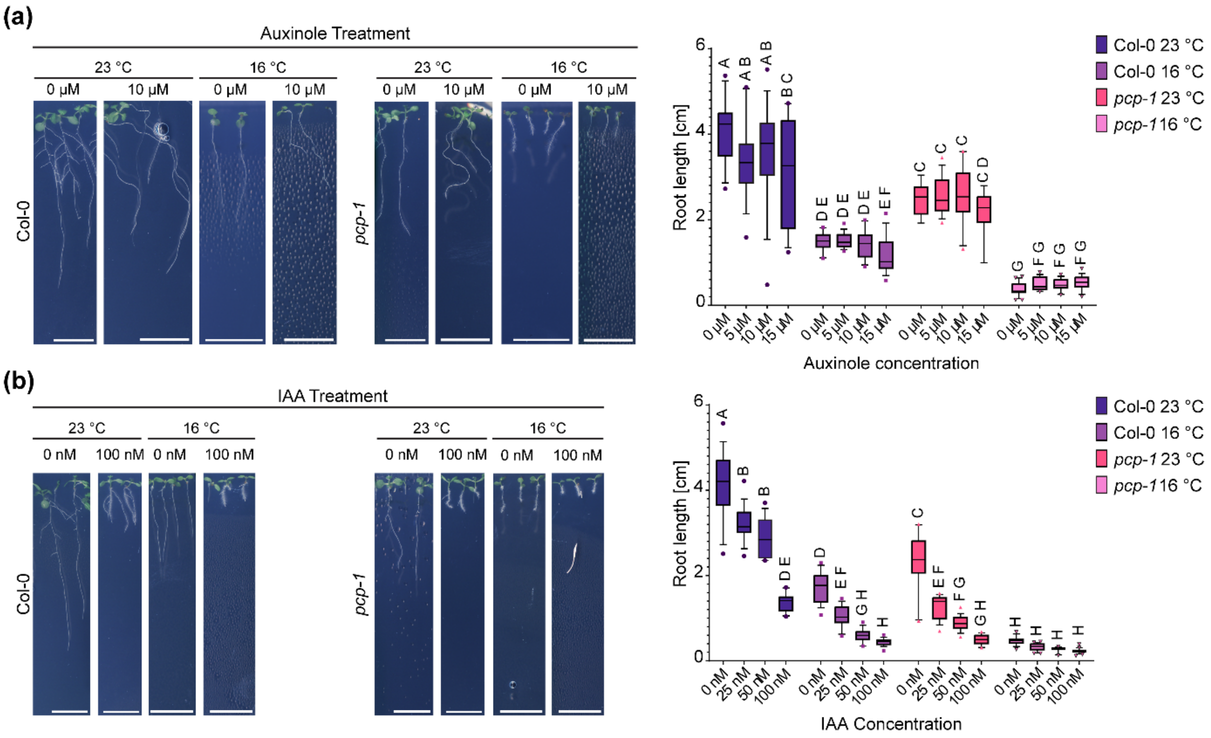


**Fig. S3 Auxinole treatment partially suppresses *pcp-1* root hair phenotype at 16°C, while IAA induces root hair phenotype in Col-0.**

(a) 10 µM auxinole treatment partially suppresses the root hair phenotype of *pcp-1* at 16°C but does not rescue root length. Scale bars: 1cm. Box plots are 10 to 90 percentile. Statistical test: Two-way ANOVA with posthoc Tukey analysis. (b) 100 nM IAA treatment at 16°C shortens Col-0 roots and induces root hair growth. *pcp-1* is hypersensitive to IAA treatment and produces shortened, hairy roots at 23°C. Scale bars: 1 cm. Box plots are 10 to 90 percentile. Statistical test: Two-way ANOVA with posthoc Tukey analysis.

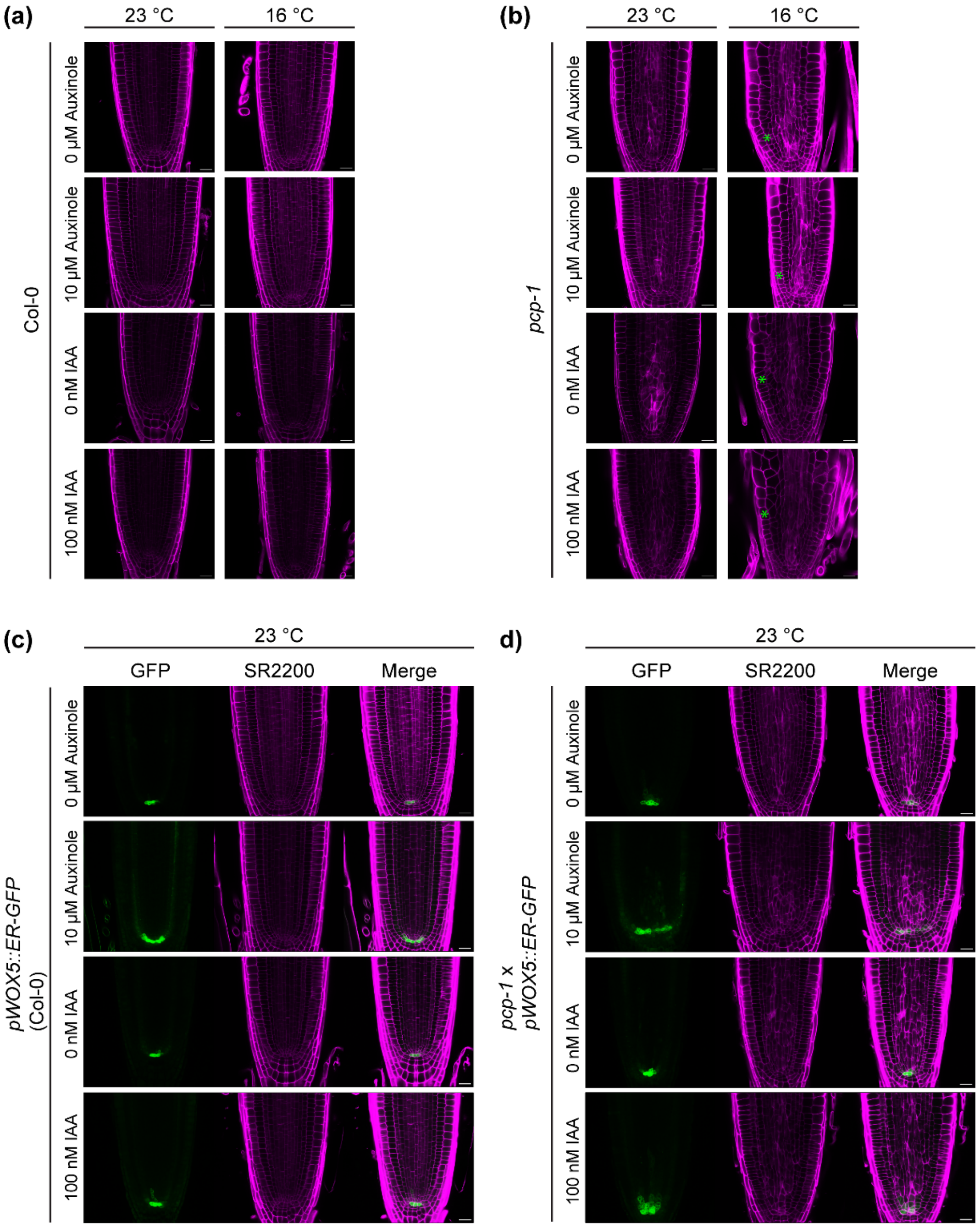


**Fig. S4 Auxinole treatment partially rescues root morphology defects in *pcp-1* at 16°C, while IAA appears to induce them.**

(a) Neither low temperature, auxinole or IAA treatment visibly alter RAM architecture in Col-0. Scale bars 20 µm. (b) RAM architecture in *pcp-1* mutants is altered at 23°C. At 16°C cortex and epidermis cells become enlarged and abnormal anticlinal and periclinal division are observed. 10 µM auxinole is sufficient to partially rescue these defects. *pcp-1* is more sensitive to IAA treatment at 23°C than Col-0, depicted by the disruption of QC morphology. At 16°C this sensitivity becomes more evident, as seen by the enlargement of epidermal, cortical, and endodermal cells, and a disruption of the QC. Green asterisks indicate positions of structural changes due to temperature or hormone treatment. Scale bars 20 µm. (c-d) At 23°C auxinole treatment enlarges the area of *pWOX5::ER-GFP* expression in both Col-0 and *pcp-1*. IAA treatment induces *pWOX5::ER-GFP* expression in stele initials in Col-0. *pcp-1* reacts more sensitively to the treatment, *pWOX5::ER-GFP* expression can be observed stretching further into the stele. Scale bars: 20 µm.

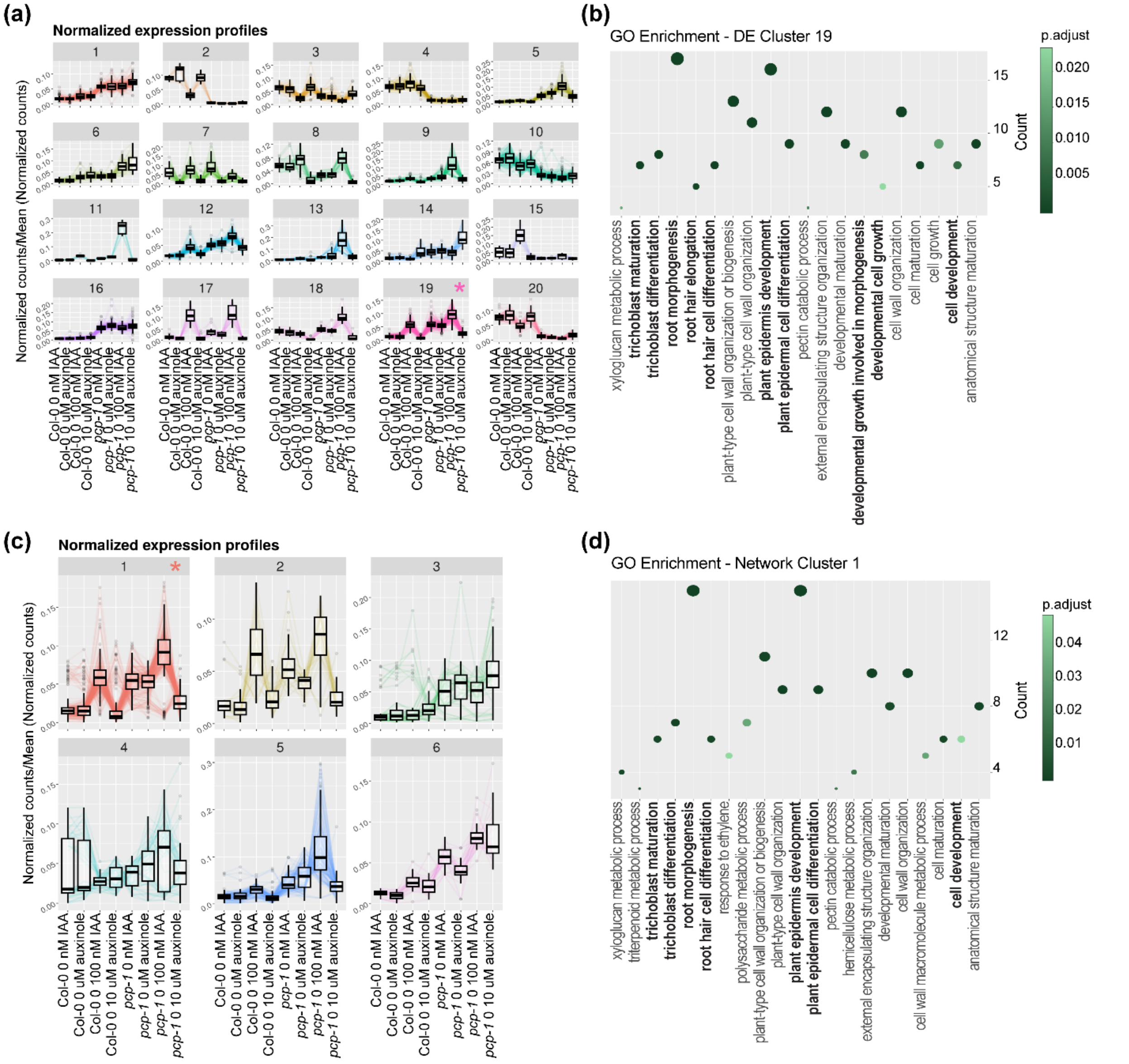


**Fig. S5 Clustering analysis and network inference of RNA sequencing data.**

(a-b) Genes in cluster 19 show an expression pattern in agreement with phenotypic observations. Gene ontology enrichment (biological process, performed in DIANE) reveals enrichment in processes linked to root hair development and root morphology. (c-d) Network inference of the sequencing data points towards cluster 1 based on expression profile. This cluster is enriched in processes involved in root hair development and root morphology. In (b) and (d), the size of the circles represents the number of genes present in the enriched GO term, and the color gradient represents the adjusted p-values.

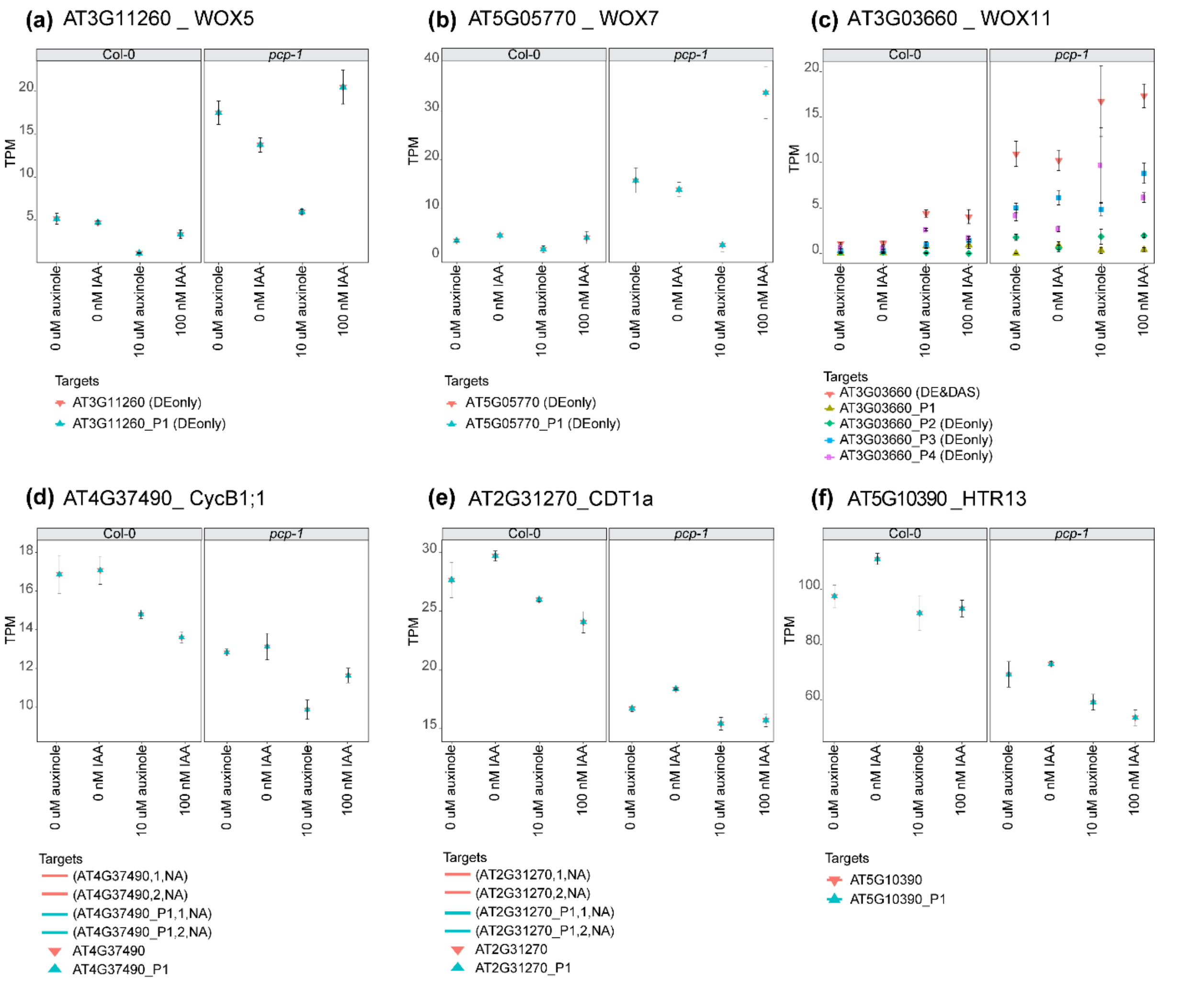


**Fig. S6 RNA sequencing expression data of meristematic regulators and cell cycle indicators.** (a-b) *WOX5* and *WOX7* expressions are relatively stable in Col-0 but are induced in *pcp-1* control conditions and in response to IAA treatment. Auxinole treatment reduces expression levels. (c) *WOX11* expression is elevated in *pcp-1*. The expression of different *WOX11* isoforms is altered in *pcp-1*. (d) *CycB1;1* expression in reduced in response to IAA and auxinole treatment in both Col-0 and *pcp-1*. Furthermore, *CycB1;1* expression is overall reduced in *pcp-1* mutants. (e-f) *CDT1a* and *HTR13* expression is reduced in *pcp-1*.

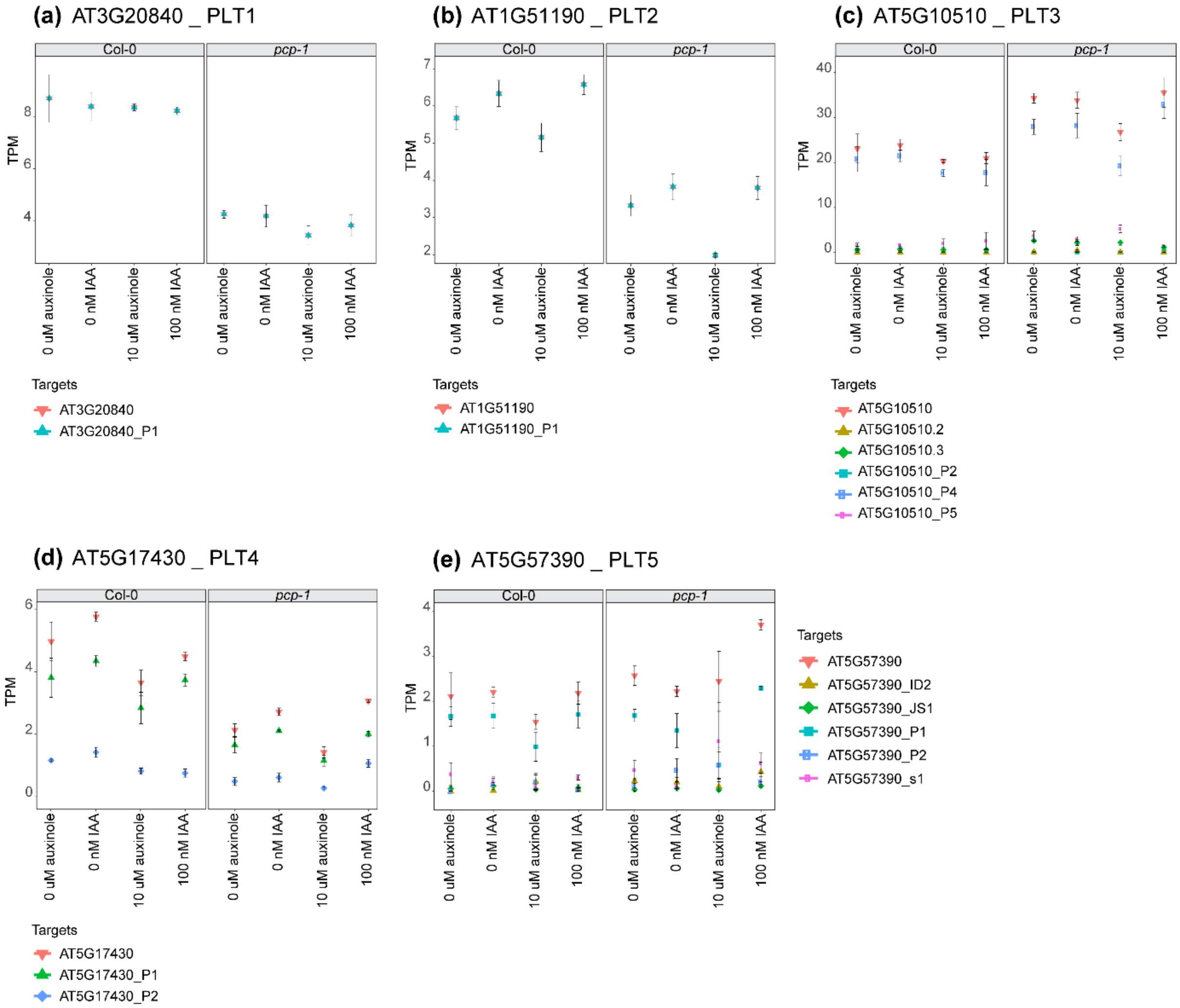


**Fig. S7 RNA sequencing expression data PLT genes.**

(a-b) *PLT1* and *PLT2* expression is reduced in *pcp-1*, without any evident treatment effect. (c) *PLT3* expression is mildly elevated in *pcp-1*. Auxinole treatment slightly lowers *PLT3* expression. (d) *PLT4* expression is relatively low in both genotypes, but slightly lower in *pcp-1*. (e) *PLT5* expression is low, but not significantly affected by treatment or genotype.

**Table S1** *Arabidopsis thaliana* plant material used in this study.

| **Plant ID** | **Accession Number** | **Selection** | **Reference** |
|---|---|---|---|
| Col-0 | NA | NA | NA |
| *pcp-1* | SALK_089521C | Genotyping | Capovilla *et al.,* 2018 |
| *pDR5::SAND* | | BastaR | Marquès-Bueno *et al.,* 2016 available at: http://www.ens-lyon.fr/RDP/SiCE/SWELLINE.html |
| *pGL2::SAND* | | BastaR | |
| *pEXP7::SAND* | | BastaR | |
| *pWOX5::ER-GFP* | | Fluorescence | Sarkar *et al.,* 2007 |
| *yuc1D* | | Phenotyping | Zhao *et al.,* 2001 |
| PlaCCI | | Fluorescence | Desvoyes *et al.,* 2020 |

**Table S2** *PIN protein immunolocalization: Antibody dilutions.*

| **Antibody** | **Dilution** | | **Source** |
|---|---|---|---|
| Anti-PIN1 (sheep) | 23°C | 1:500 | NASC ID: N782246, provided by Dr. Ranjan Swarup |
| | 16°C | 1:300 | |
| Anti-PIN2 (rabbit) | 23°C | 1:1000 | provided by Dr. Siamsa Doyle |
| | 16°C | 1:500 | |
| Anti-PIN7 (rabbit) | 23°C | 1:500 | provided by Dr. Siamsa Doyle |
| | 16°C | 1:300 | |
| CY3 anti-rabbit | 23°C | 1:400 | Jackson ImmunoResearch, 713-165-003 |
| | 16°C | 1:250 | |
| CY3 anti-sheep | 23°C | 1:200 | Jackson ImmunoResearch, 111-165-003 |
| | 16°C | 1:100 | |

**Table S3** *Confocal Microscopy settings.*

| Fluorescent marker | Excitation wavelength (nm) | Emission wavelength (nm) |
|---|---|---|
| SR2200 | 405 | 419-501 |
| SAND (4x-YFP) | 488 | 498-598 |
| GFP | 488 | 498-598 |
| CY3 | 514 | 538-586 |
| CFP | 458 | 464-507 |
| YFP | 514 | 517-589 |
| mCherry | 561 | 588-697 |

**The following tables can be found as separate excel tables:**

**Table S4** Raw data of auxin quantification and statistical analysis for each auxin compound.

**Table S5** Raw data of cytokinin quantification and statistical analysis for each cytokinin compound.

**Table S6** Statistical analysis sheets of root lengths, number of cells in proximal meristem and stele at 27°C, 23°C, 16°C and shifted.

**Table S7** Statistical analysis of WOX5 and CycB1;1 cell counts.

**Table S8** RNAseq results derived from the 3D-App, including DE genes and transcripts, DAS genes, DTU transcripts, cluster analysis and GO enrichment of cluster 12. Analysis results from DIANE, encompassing GO enrichment of cluster 19 and network data cluster 1.

**Methods S1 PIN protein immunolocalization in Arabidopsis thaliana roots using the InsituProVSI Robot**

For PIN protein immunolocalization, seedlings were fixed in 4% w/v paraformaldehyde (158127, Sigma-Aldrich) for 1h. During this step, fresh solutions of 2% w/v Driselase™ Basidiomycetes sp. (D8037, Sigma-Aldrich), 3% v/v IGEPAL® CA-630 (I8896, Sigma-Aldrich), blocking solution (3% w/v milk), 0.1% Triton™ X-100 (Merck, T8787) in PBS, 0.1% Triton™ X-100 in sterile water, and sterile water were prepared. Primary and secondary antibodies were diluted in blocking solution as follows: for seedlings grown at 23°C, we used anti-PIN1 1:500, anti- PIN2 1:1000, anti-PIN7 1:500, anti-rabbit 1:400 and anti-sheep 1:200. For seedlings grown at 16°C we used: anti-PIN1 1:300, anti-PIN2 1:500, anti-PIN7 1:300, anti-rabbit 1:250 and anti-sheep 1:100. After fixation, seedlings were transferred to InsituProVSI robot wells, containing 130 μL of 0.1% Triton™ X-100 in PBS. We followed the protocol described in Sauer et al., 2006 and Doyle et al., 2015, but briefly, samples were washed 3x in 0.1% Triton™ X-100 in PBS, and consecutively washed 3x in 0.1% Triton™ X-100 in sterile water. Samples were then treated with Driselase™ and incubated at 37°C for 30min, followed by three washes with 0.1% Triton™ X-100 in PBS. Then, samples were treated with IGEPAL and incubated at room temperature for 30min. This step was repeated twice, and then samples were washed 3x with 0.1% Triton™ X-100 in PBS. The samples were then incubated for 1 h in blocking solution, followed by three washes with 0.1% Triton™ X-100 in PBS. Subsequently, the primary antibody was added, and the samples were incubated at 37°C for 4 h, washed 3x with 0.1% Triton™ X-100 in PBS, then the secondary antibody was added, and samples were incubated at 37°C for 4 h. Finally, the samples were washed 3x with 0.1% Triton™ X-100 in PBS, and then 3x with water. The samples were then carefully transferred to microscopy slides, using glycerol as mounting medium, and immediately imaged.
